# *De Novo*, Post-Zygotic, Inter-Tissue Mosaicism of Cell Autonomous *ADNP* Mutations in Autistic Individuals: Restricted Environmental Contribution

**DOI:** 10.1101/2022.06.21.496616

**Authors:** Mohiuddin Mohiuddin, Zlatko Marusic, Mirna Anicic, Van Dijck Anke, Elisa Cappuyns, Rizzuti Ludovico, Alessandro Vitriolo, Gal Hacohen Kleiman, Iris Grigg, Giuseppe Testa, Illana Gozes, R. Frank Kooy, Christopher E. Pearson

## Abstract

Many neurodevelopmental disorders, including autism, are caused by *de novo* mutations, that might arise as early as in the parental germline, during embryonic, fetal development, or as late as post-natal aging. Intra-tissue mutation-load variations could impact clinical presentation. One of the most common causes of autism is *de novo* mutations in *ADNP*. We developed an ultra-sensitive, highly-quantitative droplet digital PCR assay to determine *ADNP* mutation levels in patient tissues, including blood, teeth, hair, and 24 different tissues from a post-mortem *de novo ADNP*-mutated child (∼6-years old), including a transplanted liver from a non-mutant donor (retained for 22 months). Striking variations of *ADNP* mosaicism arose between tissues of the same individual. Mutation load differences were evident between post-mortem tissues, but not in the transplanted liver — supporting a cell autonomous genetic vulnerability to *de novo* mutations, arguing against a transferable environmentally-sensitive DNA damage/mutation predisposition. Variations between tissues suggest a developmental timing of the mutations. Most individuals showed at least one tissue with less than heterozygous mutations, where the presence of the homozygous non-mutant cells indicates that *de novo ADNP* mutations arose post-zygotically. Highly variable *ADNP* mosaicism between tissues, that within an individual can be less than heterozygous or approach homozygosity, indicate rapid ongoing post-zygotic, and possibly post-natal, somatic mutations, contributing to clinical variability.

## INTRODUCTION

Autism spectrum disorder (ASD) is a common form of neurodevelopmental intellectual disorder characterized by a combination of deficits in social interaction and communication together with restrictive and repetitive behaviors(1). The genetic and environmental causes of autism and how this correlates with the extreme clinical variability between affected individuals is poorly understood, a knowledge that would be useful in clinical assessment, prognosis, and management.

For over a decade *de novo* mutations have been associated with ASD, in cases where affected offspring, but neither parent, show one of a series of recurrent mutations(2). In the past decade, most research efforts have focused upon the pathogenic disease mechanisms of *de novo* mutation(3, 4) and on the identification of new disease-associated *de novo* mutations. For example, *de novo* mutations are now linked to numerous neurodevelopmental and neuropsychiatric diseases, including ASD, epilepsy, schizophrenia, and cerebral cortical malformations(5). Less attention has been given to how *de novo* mutations arise or how their continued accumulation may worsen disease(6).

It is important to understand the timing, effect of ageing, and tissue-specificity of *de novo* mutations, as well as the possible contribution of environmental insult to ASD-associated *de novo* mutations(7–10). For example, as recently suggested, environmental/nutritional insults may contribute to ASD-associated *de novo* mutation, where affected individuals, through unique gene-environment interactions, may show elevated sensitivity/vulnerability to exposure-induced mutagenesis of ASD susceptibility genes events(7). *De novo* mutations may arise in the parental germline, during embryogenesis, fetal development, or postnatally. Knowing if disease-associated *de novo* mutations arise pre- or post-zygotically, and over the course of ageing, could inform on lifestyle choices to minimize exacerbating *de novo* mutations.

Somatic mutations, by definition occur post-zygotically. Recent studies on mutation loads in different tissues of unaffected human donors provided insights into cell lineage commitment from germ lineage, through embryonic, fetal, and post-natal development in different parts of the body(11–19). Surprisingly, the germ lineage had fewer mutations than somatic tissues, with evident mutation signatures induced by exogenous and endogenous mutagens(13). During the first few embryonic divisions, the somatic mutation rate is comparatively high, approximately 2.4 mutations per cell per generation. Cells establish more developed DNA-repair mechanisms from the 4-cell stage onwards, leading to a reduced mutation rate(11, 12). During the lifespan of cells, random mutations are accumulated and passed on to all their daughter cells. Thus, mutation profile of a cell can unveil important insights into the biology and evolution of tissues in later life(13, 14).

Somatic mutations can lead to tissue specific mosaicism of mutation loads, and can lead to the existence of genetically different cells within a single organism. Variable levels of somatic mosaicism have been reported in induced pluripotent stem cells and human tissues, including skin, brain, and blood(20–24), with the highest mutation load in the intestine(13).

The somatic mutation rate can be high during neurogenesis, where such mutations may lead to neurodevelopmental disease(25). Recent analyses of the human central nervous system (CNS) reveal that the CNS genome is more susceptible to high levels of somatic mutations(26). Mounting evidence indicates that somatic mutations contribute to various neurodevelopmental and neuropsychiatric disorders and neurodegenerative diseases(26–33).

Here we study *de novo* mutations in *ADNP* (OMIM#611386), which is one of the most frequently *de novo* mutated genes, being responsible for ∼0.2% to as many as 1.1% of ASD cases(34–39). *ADNP* is a *de novo* mutated gene, as determined through multiple studies of blood DNA ASD/ID cohorts where affected offspring, but neither parent shows the mutation. Helsmoortel–Van der Aa syndrome (OMIM#615873; Orphanet—https://www.orpha.net/consor/cgi-bin/OC_Exp.php?lng=EN&Expert=404448), also known as *ADNP* syndrome, is a complex developmental disorder and effects multiple organ functions(34,40,41).

The *A*ctivity-dependent *n*europrotective *p*rotein, ADNP, is a putative transcription factor that is part of the ChAHP (CHD4-ADNP-HP1) complex, where ADNP acts as a competitor of CTCF binding, providing an epigenetic role for ADNP(42–46). Consistent with this, *de novo ADNP* mutations show epigenetic dysregulation(47, 48).

The high frequency of clustered autism-associated mutations of *ADNP* at a stem-loop forming(42–46) sequence revealed mutation hotspots(34). The potential to form an unusual DNA structure may predispose these sequences to mutate during aberrant DNA replication or repair in proliferating or non-proliferating tissues. It is not known if the *de novo* mutations in *ADNP* were incurred in the parental germline and/or post-zygotically, through the development and growth of the affected individual. The latter might be expected to display some degree of somatic mosaicism of the mutant allele, the degree of which may vary between tissues, which add to the clinical variations in affected individuals(40).

Here we developed a novel assay using the ultra-sensitive droplet digital PCR (ddPCR) to quantitatively assess the levels of somatic *ADNP* mutations in DNA of *ADNP* patients, from teeth and hair roots, and from post-mortem organs, addressing hotspot mutations(49) (**Table 1**). The power of ddPCR is evidenced by the detection of cancer-associated mutations at levels as low as 0.1%(50). We have shown striking evidence for somatic mosaicism of *de novo ADNP* mutations (with a sensitivity of 0.01%), arguing for mutations incurred through development, which may contribute to varying disease manifestations.

**Table 1:** Summary of mutations for the reported patients.

| Patient No. (ID) | Mutation in genomic DNA<br>(chr. 20) | Mutation in cDNA<br>(NM_015339.2) | Mutation<br>type | Inheritance |
| --- | --- | --- | --- | --- |
| 1 (111294) | g.49508752_49508755delTTTA | c.2496_2499delTAAA | Frameshift | *De novo* |
| 6 (122793) | g.49508757_49508760delTTAA | c.2491_2494delTTAA | Frameshift | *De novo* |
| 8 (2376) | g.49508757_49508760delTTAA | c.2491_2494delTTAA | Frameshift | *De novo* |
| 15 (3314531?) | g.49508757_49508760delTTAA | c.2491_2494delTTAA | Frameshift | *De novo* |
| 24 (89686) | g.49508752_49508755delTTTA | c.2496_2499delTAAA | Frameshift | *De novo* |
| 32 (04719-11) | g.49508752_49508755delTTTA | c.2496_2499delTAAA | Frameshift | *De novo* |
| 40 (NA) | g.49508752_49508755delTTTA | c.2496_2499delTAAA | Frameshift | *De novo* |
| 100 (NA) | g.49508752_49508755delTTTA | c.2496_2499delTAAA | Frameshift | *De novo* |
| Postmortem | g.49509757duplication | C.1676dup | Frameshift | *De novo* |

## RESULTS

### Development and validation of an ultra-sensitive *ADNP* mutation load assay

The high frequency of clustered mutations of *ADNP* at the stem-loop forming sequences (Table 1) supports this region as a mutation hotspot (**Supplementary Figure 1**)(34). We hypothesize that *ADNP* may incur somatic mutations, where inter-tissue variations would be indicative of mutation accumulation. Stem-loop DNA structures may form at the *ADNP* gene during DNA replication or DNA repair and incorrect processing of these structures may lead to mutations. The *ADNP* mutation hotspot is indicated in **Supplementary Figure 1** (top): 5’ TTGAC***TTTA***<u>TTTA</u>TTCTTTCTTTC-3’, where the bolded italicized ***TTTA*** sequences are frequently deleted in patients(40). Interestingly, this TTTA, is immediately followed by another TTTA (underlined and buried within a T-rich strand) both of which could biophysically permit misaligned base-pairing forming slipped-out regions of TTTA during processes such as DNA replication or DNA repair(51). Notably, such stem-loop DNA structures have been identified to form at the mutant expanded CTG/CAG repeat tract of the *DMPK* gene in tissues of myotonic dystrophy patients, where the levels of the stem-loop structures correlated with the levels of somatic repeat instability(52). Unusual DNA structures may contribute to *de novo ADNP* mutations.

To sensitively evaluate the degree of mosaicism for *ADNP* mutation in different cells and tissues, we developed a novel assay using the ultra-sensitive droplet digital PCR (ddPCR), which is a powerful non-sequencing based method to sensitively and accurately quantify nucleic acid templates(53–60). Digital PCR improves quantification of mosaicism by segregating non-mutant from mutant alleles into individual droplets. Partitioning of a PCR reactions into ∼20,000 droplets, where sample dilutions are low-enough so that a proportion of droplets contain no template DNA molecules, and PCR amplification of a single template occurs in isolation within each individual droplet. Following amplification, droplets containing the target sequence are detected by fluorescence and scored as positive, while non-fluorescent droplets are scored as negative(54,57,59,60). Two targets can be detected simultaneously. Poisson statistical analysis of the number of positive and negative droplets yields absolute quantification of the target sequence(57, 59). Sensitivity and specificity of mosaicism are improved because each compartment, on average, contains only one type of target DNA – mutant, non-mutant, or no target. This enables precise, highly sensitive, and accurate quantification of nucleic acid variants. Sensitivity of mutation detection at some loci by ddPCR has been reported at levels as low as 0.1%(50).

Non-competing (hybrid) duplex ddPCR assays enable simultaneous amplification of two DNA targets within a single reaction(56). The two probes are each labelled with a different dye to match the two detection channels (**Figure 1A**). There are four possible configurations of amplicon products that may arise in any given ddPCR reaction, where only three of them are informative, as described below. The universal/reference probe (HEX probe, green) targets an area of the amplicon that is not expected to be variable and thus, provides a reference for the total number of molecules present in the sample, irrespective of sequence. The variant probe (FAM probe, blue) targets an area of the amplicon containing a mutable site (c.2496-2499delTAAA), binding only when the amplicon is mutated (**Figure 1A**). We observed the following cluster configurations: (i) negative droplet partitions that contain no targets for either probe, indicating an absence of DNA target template (**Figure 1B**); (ii) single-positive cluster for the universal probe (wild-type/non-mutant only droplet partitions) (**Figure 1C**); and (iii) double-positive cluster for both the universal (green) and variant (blue) probes.

**Figure 1.**
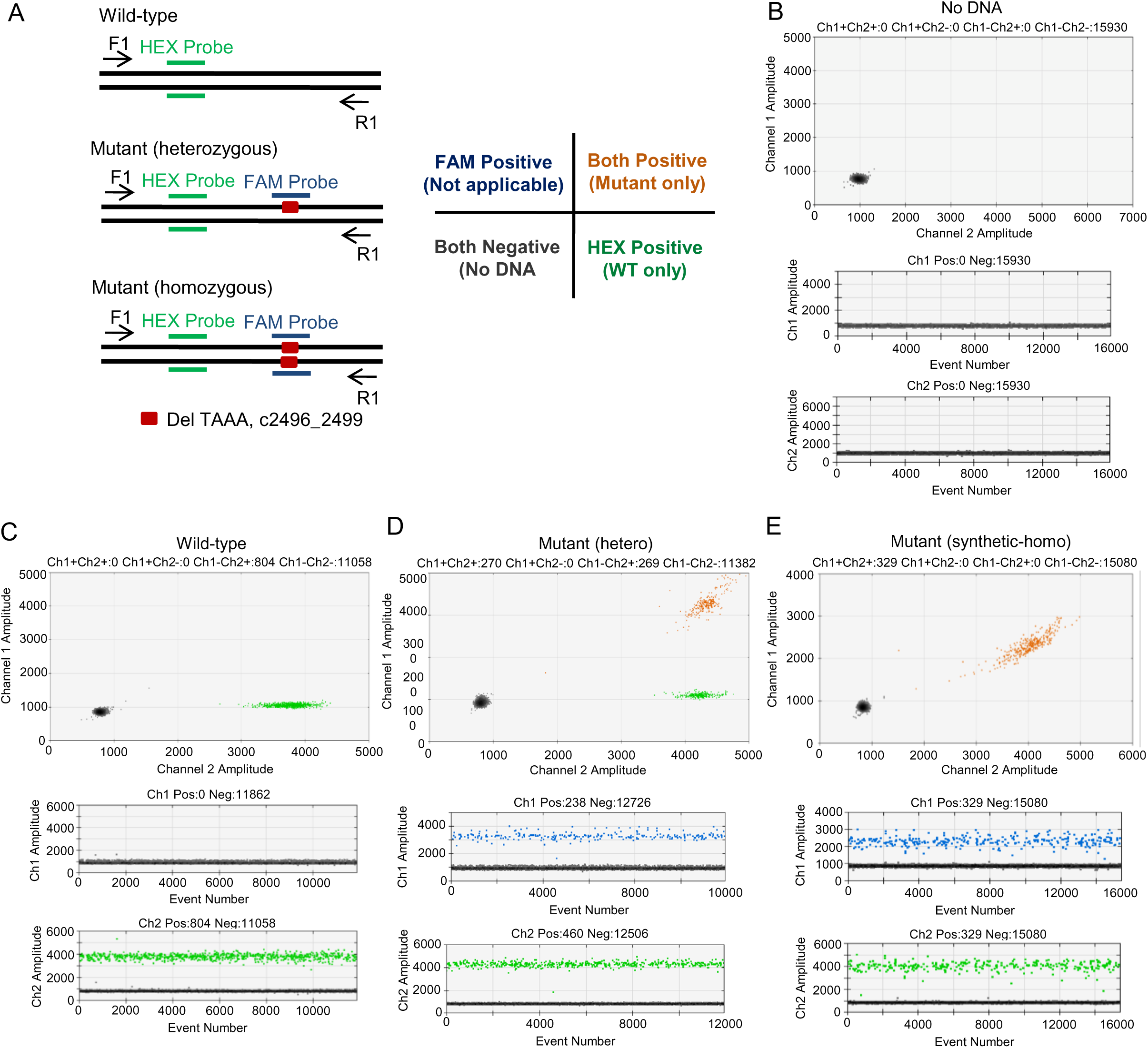
Proof-of-principle of the *ADNP* ddPCR assay. (A) Schematic illustration for non-competing (hybrid) duplex reactions of ddPCR assay to distinguish wild-type and mutant allele of *ADNP* gene (left panel). HEX probe (universal / reference probe) is designed upstream of mutation site (c.2496_2499TAAAdel), whereas FAM probe is specific for mutation region. Primer pair (F1/R1) amplifies a sequence covering both HEX and FAM probes specific sequences. Only three clusters are visible in the non-competing (hybrid) duplex reactions (right panel). (B-E) Upper panel: 2-D plot showing amplification of *ADNP* gene in the Blank (B), Wild-type (C), Heterozygous mutant (D) and Synthetic homozygous mutant (E) DNAs. Lower Panel: 1-D plot showing *ADNP* gene amplification. A well-defined separation between negative droplets and positive droplets for both FAM and HEX probes was seen. Wild-type and mutant allele of *ADNP* gene can be distinguished by ddPCR.

The double positive reads as cyan, a mixture of the green and blue probes. The missing single-positive cluster (variant only) is embedded into the double-positive clusters since the universal probe always produces a signal in the presence of any amplicon (**Figure 1D**). We simulated the fully-homozygous mutant state by using a cloned *ADNP* amplicon template fragment, that had been confirmed by sequencing to have the mutant allele. For this synthetic homozygous mutant, wild-type/non-mutant only partition is not present (**Figure 1E**), providing proof-of-principle that the ddPCR assay can distinguish wild-type and mutant *ADNP* alleles. The term “absolute quantification” used in ddPCR, refers to an estimate derived from the count of the proportion of positive partitions relative to the total number of partitions and their known volume(57). This subdivision enables quantification to be performed with high precision, independent of a standard curve (**Supplementary Figure 2**)(58).

The ddPCR is an end-point assay with the determination of the positive droplet fraction and Poisson statistics calculating the absolute number of starting copies making calibration curves unnecessary(55, 59). Due to the simultaneous presence of reference amplification in each reaction during ddPCR, as well as a consequence of unprecedented precision, replicates are not needed in cases with target concentrations above the limit of detection. ddPCR is highly suitable for the detection of rare mutation events, even in the presence of high background. Using a similar approach to Mika *et al*(53), our quality control test on artificial *ADNP* mosaicism mixture samples revealed a high correlation between the estimated and the observed percentage values for two discriminating markers. The limit of detection of *ADNP* mutations by ddPCR was estimated to be as low as 0.01% (**Supplementary Figure 2D**) permitting a sensitive and reliable determination of mosaicism. Additionally, ddPCR is a direct quantitative technique that does not rely on an external standard/reference curve, giving higher quantitative accuracy and precision(57). To test this strength, we generated a synthetic fully-homozygous mutant (c.2496-2499delTAAA) DNA from *ADNP* patient DNA (heterozygous mutant) (**Supplementary Figure 2A**) and mixed this with non-mutant (wild-type) DNA in different ratios: 0, 0.2, 0.4, 0.8, 1.2, 1.6 and 2.0 (mutant/wild-type). The number of mutant and wild-type alleles were quantified by ddPCR. Based upon the ratio of mutant allele/wild-type allele, a standard curve (linear regression) was calculated. Mixed ratios of DNAs (x-axis) and the ratio of FAM and HEX positive droplets (y-axis) were almost the same (**Supplementary Figure 2B-C)**, indicating that ddPCR allows quantifying samples without using a standard curve.

### Degree of mosaicism for *ADNP* mutation in different tissues of *ADNP*-patients

Using our sensitive *ADNP* mutation assay we first assessed the degree of mosaicism for the above noted *ADNP* c.2496-2499TAAAdel mutation (**Supplementary Figure 3)** in DNA extracted from a range of *ADNP* patient tissues from multiple patients, including blood, teeth, hair root cells (ectodermal origin), and patient derived lymphoblastoid cell lines. A 1:1 ratio of wild-type versus mutant allele was reflective of the heterozygous mutant state. Results showed high levels of somatic mosaicism of *ADNP* mutations between tissues of the same individual: in one case *ADNP* mutation loads approached homozygosity in the hair (allelic ratio of wild-type versus mutant allele was greater than one) but were less than heterozygosity in the teeth (allelic ratio of wild-type versus mutant allele was less than one) (**Figure 2 and Supplementary Figure 4**), indicating that this individual is mosaic for the c.2496-2499TAAAdel *ADNP* mutation. That a less that heterozygous state was detected indicates that some cells in the population were in fact homozygous for the non-mutant *ADNP*. We also assessed the degree of mosaicism for the c.2491-2494TTAAdel *ADNP* mutation (**Figure 2 and Supplementary Figures 5, 6)** in DNA extracted from a range of *ADNP* patient-derived lymphoblastoid cell lines. Varying levels of mosaicism were also evident. It should be noted that de novo mutations may be introduced and propagated during lymphoblastoid cell line transformation that are unrelated to disease biology(61, 62). To investigate possible cross-contamination, we applied the c.2496-2499TAAAdel mutation specific probes to DNAs of *ADNP* patients devoid of that mutation (patient with a mutation c.1676dup A). We could not detect any positive droplets, indicating that there is no cross-contamination (**Supplementary Figure 7**). Moreover, as expected, tissues from the parents did not show *ADNP* mutations (**Figure 2D**).

**Figure 2.**
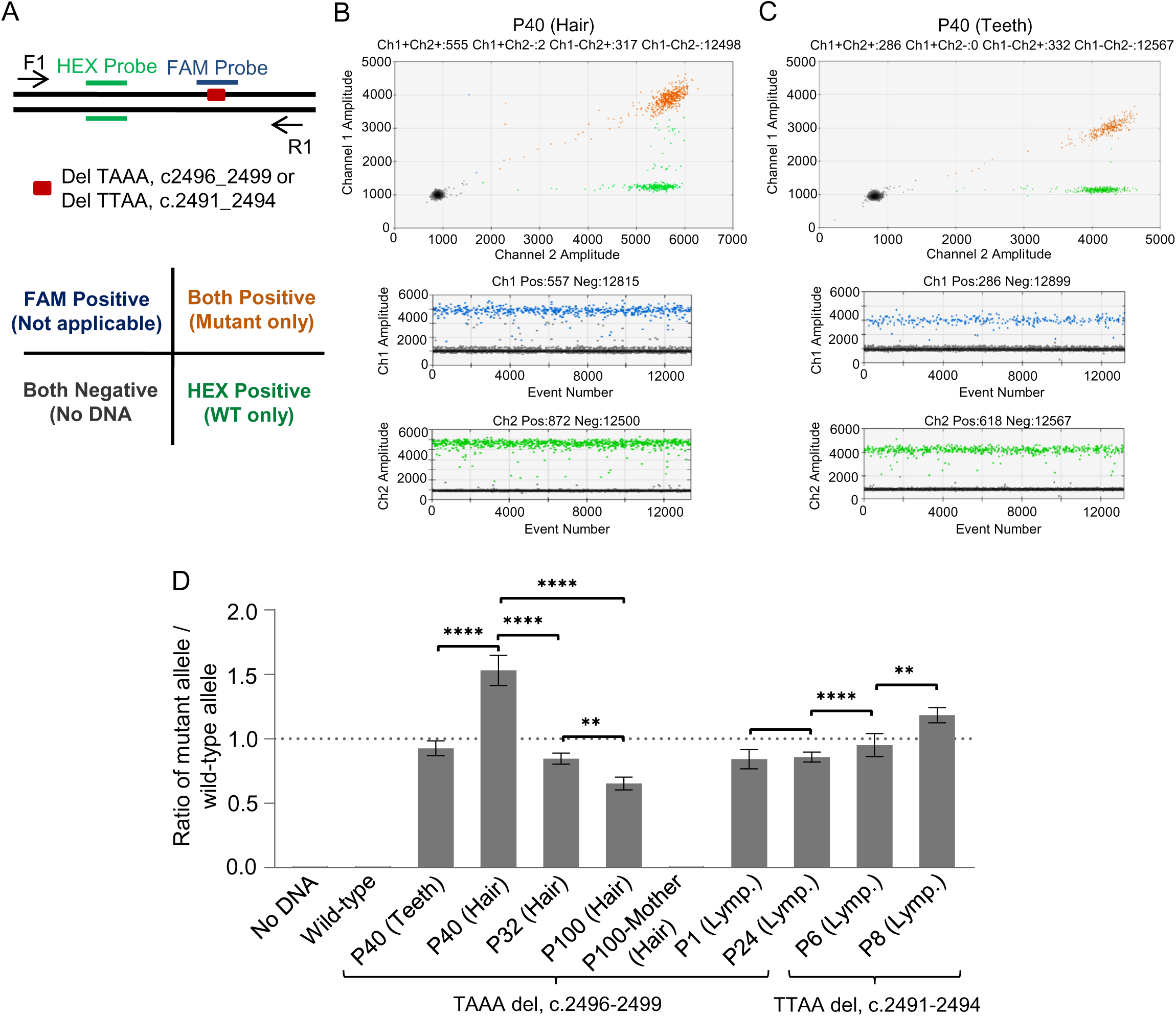
Degree of mosaicism for *ADNP* mutation (c.2496_2499TAAAdel or c.2491_2494TTAAdel) in different tissues, and lymphoblastoid cells. (A) Schematic illustration for non-competing (hybrid) duplex reactions of ddPCR assay to evaluate mutation load in different tissues, and lymphoblastoid cells (upper panel). HEX probe (universal / reference probe) is designed upstream of mutation site, whereas FAM probe is specific for mutation region (c.2496_2499TAAAdel or c.2491_2494TTAAdel). Primer pair (F1/R1) amplifies a sequence covering both HEX and FAM probes specific sequences. Only three clusters are visible in the non-competing (hybrid) duplex reactions (lower panel). (B-C) Upper panel: 2-D plot showing amplification of *ADNP* gene in the P40-hair (B) and P40-teeth (C) DNAs. Lower Panel: 1-D plot showing *ADNP* gene amplification. A well-defined separation between negative droplets and positive droplets for both FAM and HEX probes was seen. (D) Histogram representing the ratio of mutant vs wild-type allele (*y*-axis) in the indicated cell lines and tissues derived genomic DNAs (*x*-axis). Error bars indicate SD of more than three independent experiments. Dotted line indicates the ratio for heterozygous mutation.

### Inter-tissue variations of *ADNP* mutation load in post-mortem tissues of an ASD patient

A male toddler was diagnosed with ASD and identified to have Helsmoortel–Van der Aa syndrome by sequencing of their blood DNA to have a *de novo* mutation in *ADNP* (c.1676dup). The patient was apparently heterozygous for a *de novo* mutation, that was absent in both of his parents blood DNA. Since the child had several features [premature birth (32-week gestation versus a healthy average of 38-42 weeks), unusual facial characteristics, neurodevelopmental delay, autistic behavior, severe speech delay, seizures and liver failure of undetermined cause], whole genome sequencing was performed. Following liver failure, at ∼1 year 5-months age, the child, received a split liver transplant (the left liver lobe of a cadaver) and a liver re-transplantation at ∼3 years 5-months age (the right liver lobe of a cadaver). 22 months after the second liver transplant, the child suffered septic shock, pneumonia (*Candida lusitanie*), Epstein Barr virus infection, and succumbed to clinical complications at ∼6 years 2-months of age. An autopsy was performed and different tissue samples were taken for analysis. All tissues, but the liver, were confirmed to be from the same individual, as indicated by forensic analysis (**Supplementary Table 2**). To determine if there is tissue-specific variations in the *ADNP* mutation load, we extracted DNA from the 24 tissues listed in **Table 2**. DNA from the cerebellum and blood were Sanger sequenced across *ADNP* to validate the mutation (**Supplementary Figure 8**). We designed a FAM probe that detects the mutated c.1676dup *ADNP* sequence of this patient, whereas the HEX probe detects the wild-type *ADNP* sequence.

**Table 2:**
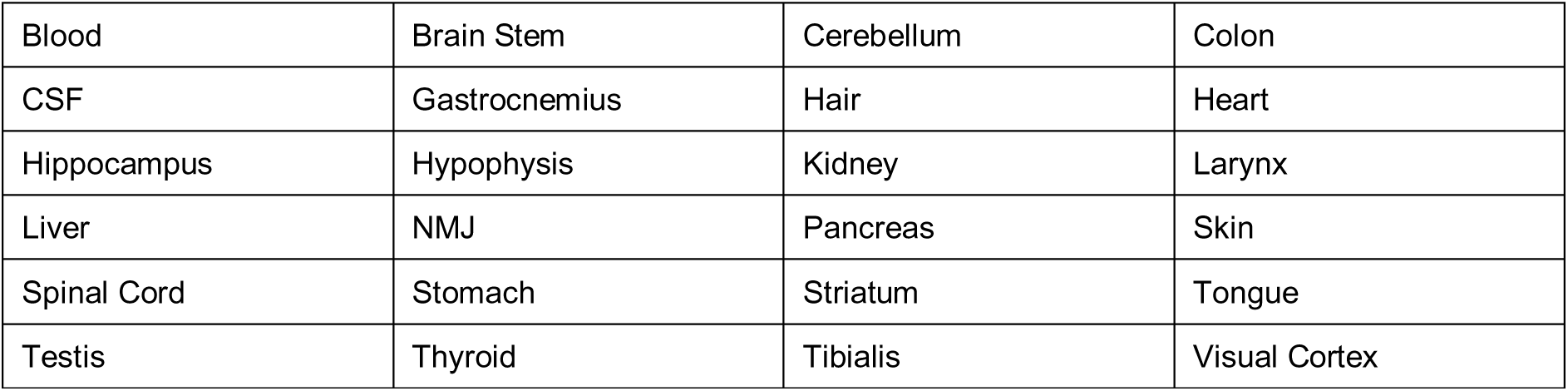
Tissues from the *ADNP* c.1676dup autopsy.

To quantify small insertion or deletion events (**Figure 3 and Supplementary Figure 9**), we applied competing duplex ddPCR assay (one primer pair with two probes binding the same region)(63–66) (**Figure 3A**). With this assay we expected to obtain four clusters of droplets. The negative partitions contain no targets for either probe (**Figure 3B**). The single-positive clusters for the universal (HEX) probe and the variant (FAM) probe represent non-mutant only partitions and mutant only partitions, respectively. The double-positive cluster, which forms an arc conformation that spans the two single-positive clusters (**Figure 3C-F and Supplementary Figure 9**), is primarily caused by partition specific competition(56). Results from ddPCR analyses showed inter-tissue variations of *ADNP* mutation loads between the post-mortem tissues of this patient. Importantly, mutations were above and below heterozygosity, but not evident in the transplanted liver (**Figure 3G**). The allelic ratio of wild-type versus mutant allele in the transplanted liver was just greater than null (**Figure 3G**).

**Figure 3.**
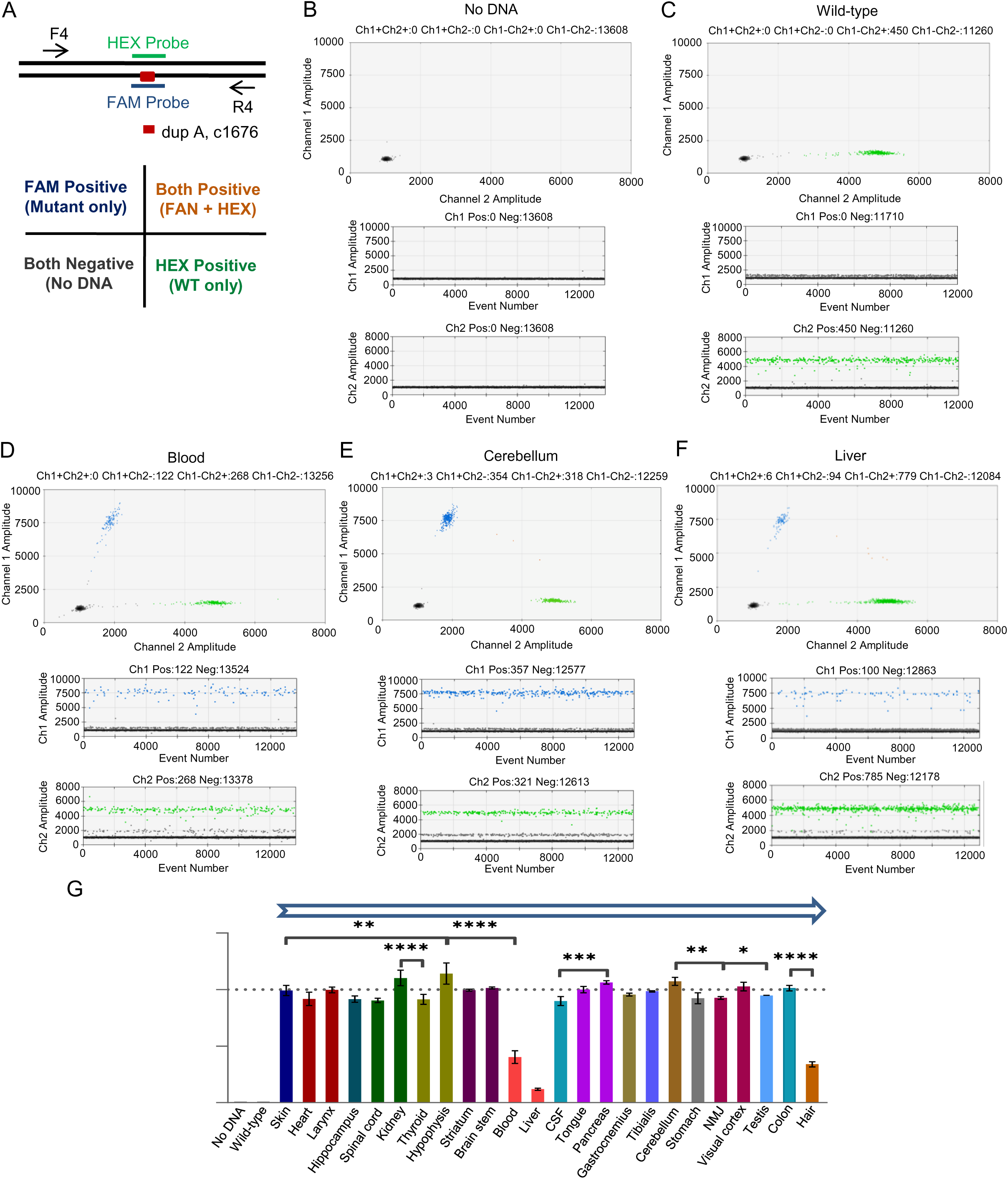
Inter-tissue variation of *ADNP* mutation loads (c.1676dup A) in tissues from a deceased *ADNP* patient. (A) Schematic illustration for competing duplex reactions of ddPCR assay to evaluate mutation loads in the post-mortem tissues of a deceased *ADNP* patient (upper panel). HEX probe (universal / reference probe) is designed for the abundant or wild-type sequence, whereas FAM probe is designed for the mutant sequence (c.1676dup A). Primer pair (F4/R4) amplifies a sequence covering c.1676 of *ADNP* gene. Unlike non-competing (hybrid) duplex reactions, four clusters are visible in the competing duplex reactions (lower panel). For the analysis of probe-competing duplex reactions to quantify a single base substitution, cross-hybridization of the probes caused by either mismatched of the probes or filter bleed-through can be visualized as a ‘leaning’ (blue) or ‘lifting’ (green) of the single positive clusters. (B-F) Upper panel: 2-D plot showing amplification of *ADNP* gene in the Blank (B), Wild-type (C), Blood (D), Cerebellum (E), and Liver (F) DNAs. Lower Panel: 1-D plot showing *ADNP* gene amplification. A well-defined separation between negative droplets and positive droplets for both FAM and HEX probes was seen. (G) Histogram representing the ratio of mutant vs wild-type allele (*y*-axis) in the indicated tissues derived from a deceased *ADNP* patient (*x*-axis). Error bars indicate SD of three independent experiments. Dotted line indicates the ratio for heterozygous mutation.

The low *ADNP* mutation load in the transplanted liver is likely due to systemic chimerism of the transplanted liver with the *ADNP*-patients’ own *ADNP*-mutated macrophage/Kupffer and blood cells(67). Transplanted livers are rapidly repopulated by the recipient’s macrophage and blood cells(67). Thus, the absence of detectable *ADNP* mutations in the transplanted liver serves as an internal control, further validating the utility of the ddPCR assay to assess *ADNP* mutation loads.

## DISCUSSION

The aim of this study was to evaluate the level of somatic *ADNP* mutation mosaicism in a collection of *ADNP* patient tissues, as this may provide insight into the origin and timing of *de novo* mutations, which could impact the development of Helsmoortel–Van der Aa syndrome. We analyzed samples from several autistic patients with various mutations in *ADNP* gene (**Table 1**). To date, due to the rarity of ASD post-mortem tissues, nearly all studies on somatic mosaicism in autism have focused upon blood DNA (mesodermal lineage) rather than of ectodermal lineage tissues, which gives rise to the brain(6). We developed an ultra-sensitive droplet digital PCR assay to assess *ADNP* mosaicism in a broad-range of *ADNP* patient-derived tissues from multiple patients, including blood, teeth, hair root cells (ectodermal/neural crest), and 24 tissues from a post-mortem *de novo ADNP* mutated child, including the transplanted liver received from a non-mutant donor (**Table 2**). Levels of mosaicism of *ADNP* mutations between tissues of the same individual (**Figure 2 and Supplementary Figures 4, 6**) and an extensive array of various post-mortem tissues (**Figure 3, and Supplementary Fig. 9**) suggests that the *ADNP* mutation load in tissues depends on mutation timing during differentiation and postnatal life(68–71). Degrees of mosaicism of *ADNP* mutations between tissues could reveal mutation timing and may shed light on disease etiology and clinical variability.

Mutations that are inherited can present constitutionally in the parent (or parents) and in all tissues of the affected offspring. In contrast, *de novo* mutations are undetectable in either parent of an affected individual, at least in the tissue typically assessed, such as blood. Mutations that greatly increase the risk of neurodevelopmental, neuropsychiatric and pediatric disorders – even as heterozygous mutations, appear to arise *de novo*. *De novo* mutations have long been presumed to be due to new mutations arising in the germline of the parents(4).

Disease-causing *de novo* mutation events can arise as early as in the parental germline, during embryonic, or fetal development, or as late as post-natally. A *de novo* germline mutation arises during gametogenesis in one of the parents and is not detectable in the parent, while it is detectable in all tissues of the child. Studies have postulated that germline mutations can be influenced by both paternal and maternal age, and these age related changes are not distributed uniformly across the genome(72, 73). Previously, it was presumed that an alternate allele frequency of ≤ 40% supports a post-zygotic mutation event rather than mutations in the parental gametes(74). Paternal sperm DNA analysis allows to identify male germline mosaicism, which may assist in predicting recurrent risk of *de novo* genetic variants associated with autism(75–77). Every individual’s genome carries approximately one *de novo* germline mutation in the exome, the protein-coding region of the genome(78). Even though every cell in the individual carries the mutation, the predominant effects of the mutation depend on the distribution of gene expression. A second-hit somatic mutations increase mosaicism(79), which approach homozygosity (**Fig. 4A**). An early or late post-zygotic mutation results in a mutation present in most or all tissues of the organism (including the leukocytes, which are generally assayed for clinical genetic testing) but in a mosaic fashion, with only a portion of all cells in each tissue harboring the mutation. During development through aging, mosaicism approaching heterozygosity (**Fig. 4B**) – indicating that timing of mutations could lead to mosaicism of less than heterozygosity to approaching homozygosity. Developmental timing of somatic mutation acquirement regulates the distribution in the whole body.

**Figure 4:**
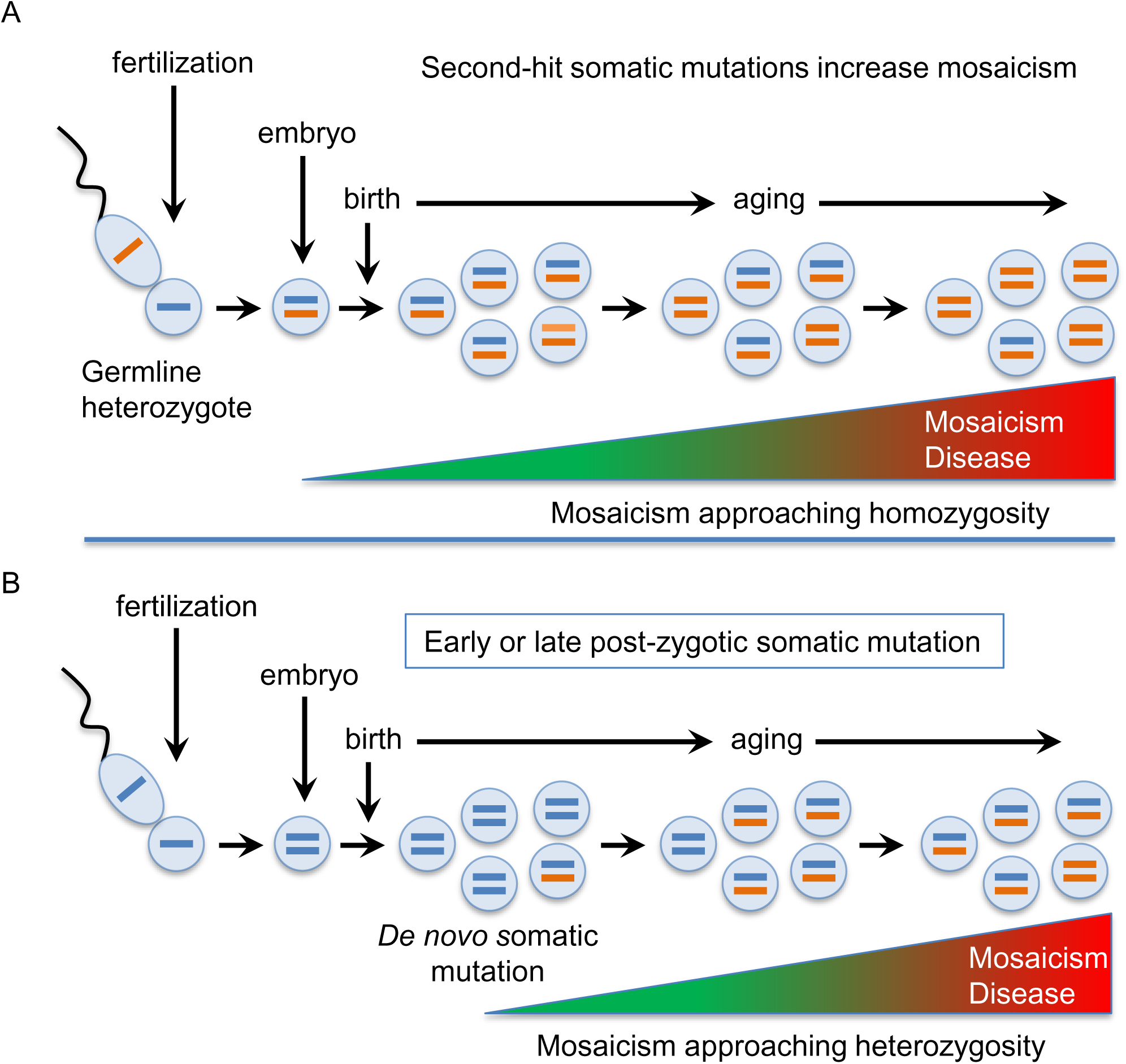
Timing of *de novo* & somatic mutations that lead to mosaicism. (A) A *de novo* germline mutation is not detectable in the parent and is detectable in all tissues of the child. Somatic mutations increase mosaicism, to approach homozygosity. (B) Post-zygotic mutation results in a mutation present in most or all tissues of the organism but in a mosaic fashion, with only a portion of all cells in each tissue harboring the mutation. During development through aging, mosaicism approaching heterozygosity. (C) Same as B, except the rate of post-zygotic somatic mutation is faster, which can approach homozygosity.

A *de novo* mutation in the parental germ line would lead to offspring with a minimum of a heterozygous mutation load (**Figure 4A**). Since most *ADNP* individuals showed at least one tissue with less than heterozygous mutations, the presence of the homozygous non-mutant cells indicates that *de novo ADNP* mutations arose post-zygotically. Since within an individual some tissues showed less than heterozygous (**Figure 4B**) and other tissues that approached homozygosity (**Figure 4C**), indicate that this are variable rates of ongoing post-zygotic somatic mutations, which may contribute to clinical variability.

The presence of the homozygous non-mutant cells indicates that *de novo ADNP* mutations arose post-zygotically. Donor-derived chimerism, the migration of donor stem cells into recipient’s tissues(67), could appear similar to mosaicism, and may lead to underestimates of ADNP mutation loads. Tissue identity was confirmed for all tissues using a forensic panel of polymorphic markers (Supplementary Table 2).

Our finding that a transplanted non-mutant *ADNP* liver, was devoid of somatic *ADNP* mutations is worthy of some comment. Liver is an organ exposed to high levels of genotoxic stress. The liver metabolizes nutrients, drugs, hormones, and metabolic waste products. Livers of non-diseased individuals can incur high frequencies of *de novo* mutations, over the course of months, detectable in young individuals (5- months)(80). The liver is particularly prone to mutations by environmental insults, with its high metabolic activity and its role in detoxification of xenobiotics(80, 81). For example, dietary choices can affect mutations in liver(82, 83). In humans, accumulation of *de novo* mutations can arise in differentiated hepatocytes with high spontaneous mutation frequencies in that significantly increase with age(80, 84). Thus, the transplanted liver was subjected to the same exposures over the last quarter of the child’s post-zygotic life (22-months), as all other organs. As a toddler, their blood revealed *ADNP* mutations, which our results suggest occurred post-zygotically. If one imagines that *de novo* somatic *ADNP* mutations continued to accumulate during the last years of the child’s life, since the transplanted liver was essentially devoid of *de novo ADNP* mutations, would argue that it is unlikely that an environmental contribution alone is responsible to the *de novo* mutations incurred by other tissues. Genetic predisposition to xenobiotically-induced *de novo ADNP* mutations by this child’s tissues is possible, as hypothesized for post-zygotic *de novo* mutations associated with ASD and other neurodevelopmental diseases(7, 8). However, a passive transmission of this predisposition to somatically mutate is unlikely, as we were unable to detect any mutations in the transplanted liver. This suggest that the *de novo* somatic *ADNP* mutations in this child were cell autonomous. The possible contribution of environmental aspects cannot be excluded(7). Future studies on the contribution of environmental insult to ongoing *de novo* ASD-associated mutations must consider the timing of the mutations, as well as a possible that may contribute to such ASD-associated *de novo* mutations(7, 8).

Cell autonomous mutation predispositions of transplanted tissues has been previously observed. Bone marrow transplant (BMT) recipients incur greatly reduced programmed somatic hypermutations in rearranged immunoglobulin V_H_ genes, leading to low memory B-cell counts compared to healthy subjects(85). It was postulated that there is a deficit in the capacity of BMT-recipient B cells to respond to signals to activate the somatic hypermutation program. The transplanted liver, in the ASD boy, at least over the 22-months that it was retained, did not incur somatic *ADNP* mutations. Non-cell autonomous effects of cells mutated in the epigenetic regulator MeCP2, have been observed. For example, *MeCP2*-mutant neurons can detrimentally affect the development of surrounding non-mutant neurons, in a non-cell-autonomous manner(86, 87). Whether disease attributes or a vulnerability to mutate present in *ADNP*-mutated cells can affect the state of surrounding non-mutant cells, is of particular interest with an appreciation of *ADNP* mutation mosaicism in a given individual. It is important to discern if ADNP traits can be laterally transferred in a non-cell autonomous manner or whether these are retained in autonomously, and whether these are affected by environmental exposure. Appreciating such factors will the illuminate that variable disease presentation(34,40,41).

We present a sensitive analysis of mutation load in an extensive collection of tissues. A previous study of various tissues of clinically unremarkable individuals for post-zygotic mutation loads revealed that different lineage trees of mutations can lead to a path(88–90).

Somatic DNA sequence variation can be acquired during the lifetime of individuals. For the disease-causing early somatic mutation, a somatic variant that arises early in embryonic or fetal development, can result in a large percentage of cells carrying a detrimental variant. Thus, mutation timing in clinical genetics should be considered seriously. The time when disease-causing *de novo* mutations continue to accumulate as somatic mutations, shifting the proportion of mutant cells within a given tissue is likely to contribute significantly to the phenotype.

Our data showed that somatic mosaicism of *ADNP* mutations arose between tissues of the same individual: in some cases, *ADNP* mutation loads were greater than heterozygosity (approaching homozygosity). Importantly, tissue specific *ADNP* mutation load differences were found between the post-mortem tissues, both above and below heterozygosity. Most individuals showed at least one tissue displaying less than heterozygous mutations, which argues against mutations having arisen in the transmitting germline and favor either age-acquired somatic mutations. The highly variable *ADNP* mosaicism loads between tissues, that can approach homozygosity, indicate extremely high rates of post-zygotic and possibly post-natal somatic mutations, and ongoing mutations may contribute to clinical variation. These mutations arise after fertilization leading to the coexistence of two or more cell populations within the same individual that can contribute to varying disease manifestations.

## Methods and Materials

### Lymphoblastoid cells, hair and teeth

After informed consent from the mother/parents/caregivers of Helsmoortel–Van der Aa syndrome children, lymphoblstoid cells and other tissues were collected.

### Postmortem collection

After informed consent from the mother (caregiver of the deceased child), postmortem tissues were collected during autopsy and documented with half undergoing immediate freezing (-1800 C) and half fixation for further histochemical analyses. Specifically, tissue samples were surgically removed and frozen in vials (brain stem, cerebellum, hippocampus, pituitary (hypophysis), striatum, visual cortex, blood, colon, muscle (gastrocnemius, tibialis and tongue), heart, kidney, pancreas, skin, spinal cord, stomach, testis and thyroid. Each tissue was homogenized using Bullet Blender® (Next Advance, Inc., NY) and the appropriate beads. A DNA sample from the cerebellum was Sanger sequenced to validate the mutation. Forensic analysis of several commonly used markers was done to confirm all tissues, but the liver, came from the same individual.

### DNA extraction

Tissues were homogenized using a Bullet Blender (Next Advance, Inc., NY). DNA were extracted from the same tissue with a ZR-Duet DNA-RNA MiniPrep Plus kit (Zymo Research, Irvine, CA). Full written informed consent was obtained from tissue donors or their relatives where appropriate.

### Digital droplet PCR (ddPCR)

To analyze somatic mosaicism, 20μl ddPCR reactions were prepared according to manufactureŕs instructions (Bio-Rad Laboratories Ltd.): 10μl of ddPCR™ Supermix for Probes (No dUTP), 900 nM each of the forward and reverse primers, 250 nM each of the fluorescence probes, 1 μL DNA (10 ng/μL), and RNase/DNase-free water. Two targets were detected simultaneously with different fluorescence probes: the FAM probe detected the mutated sequence (*ADNP*: c.2496-2499delTAAA), or in separate reactions (*ADNP*: c.2491-2494delTTAA), while the HEX probes detected the wild-type sequences. For the organ samples obtained from a deceased *ADNP* patients, the FAM probe detected the mutated (c.1676dup) sequence whereas HEX probe detected the wild-type sequence. The annealing temperature for different sets of primers and probes was established by previous gradient runs. After droplet generation, PCR was performed in C1000 Touch Thermal Cycler as follows, 95°C 10 min, 40 cycles of 94°C 30s, 58°C 1 min (for c.1211C>A mutation, 68°C 1 min; for c1676dupA mutation, 65.8°C 1 min), 98°C 10 min. Droplet analysis was performed using the QX200 instrument, and data were analyzed with Quantasoft software, with the absolute quantification (ABS) mode according to the digital MIQE guidelines. For each ddPCR sample, the same process was performed in triplicate. Data analysis was performed when the number of droplets produced was more than 10,000.

### Standard curve for ddPCR

Exon 5 of *ADNP* was amplified by PCR with primers (F5/R5) using the *ADNP* patient’s DNA as a template. PCR products were then cloned using a TOPO-TA cloning kit (Invitrogen, Carlsbad, CA). Wild type and mutant (c.2496-2499delTAAA) clones were verified by sequencing (**Supplementary Fig. 3A**). DNA was extracted from a wild type clone and a mutant using a QIAGEN Plasmid Midi Kit (Qiagen, Tokyo, Japan) and used as standard DNA to determine ratios of mosaicism. Wild-type and mutant clone DNAs were mixed in different ratios: 0, 0.2, 0.4, 0.8, 1.2, 1.6 and 2.0 (mutant/wild-type) (**Supplementary Fig. 3B)**. The number of mutant and wild-type alleles were quantified by ddPCR. Based upon the ratio of mutant allele / wild-type allele, a standard curve (linear regression) was calculated. Mixed ratios of DNAs (*x-axis*) and the ratio of FAM and HEX positive droplets (*y-axis*) were almost the same (**Supplementary Fig. 3C-D)**, indicating that ddPCR allows to quantify samples without using a standard curve.

## ACKNOWLEDGEMENTS

We thank Marc Bühler, Friedrich Miescher Institute for Biomedical Research, Basel, Switzerland for discussions, sharing unpublished results and comments on the manuscript. We thank the families for tissue sample donations.

## Author contributions

M.M. and C.E.P. conceived and designed the study. M.M. designed the ddPCR, designed and performed all experiments, processed/analyzed data, prepared figures, and wrote the original draft of the manuscript. M.M. and C.E.P. reviewed and edited the manuscript. C.E.P. supervised the study. I.G*., G.H-K., I.G^†^., V.D.A., R.L., A.V., E.C., Z.M., M.A. were involved in tissue collections/purified DNA. G.H-K., I.G., R.L., A.V., G.T., and R.F.K. critically edited the manuscript. All authors provided feedback and agreed on the final manuscript. I.G.* = Iris Grigg; I.G.^†^ = Illana Gozes.

## Methods and Materials

### Autopsy Report

Our patient was born on October 22^nd^, 2010 after the third pregnancy (first two ended as spontaneous abortions), from healthy, nonconsanguinous parents. Delivery was spontaneous vaginal in 32nd week of gestation because of cervical insufficiency, his birth weight was 1790 g, Apgar score 10/10. He was treated with phototherapy for 8 days due to neonatal hyperbilirubinaemia. He had intracranial hemorrhage gr.III. He underwent neurodevelopmental therapy because of prematurity and psychomotor delay (he started walking at the age of 1 year and 9 months and he never spoke, but did understand through gestures. He liked to play on an electronic tablet.

At the age of 1 year and 7 months, he was referred as an outpatient to our Unit for inherited metabolic diseases because of psychomotor delay. At the time, larger neurocranium, dismorphic facial features (broad forehead, flat nasal bridge, abnormally shaped ears), and palpable liver (4 cm) were observed. Laboratory tests (aminotransferases, electrolytes, organic acids) and fundus were normal.

At the age of 2 years and six months, a few days after he had an upper respiratory tract infection, he became jaundiced, had pale stools and dark brown urine, and was admitted to hospital. Laboratory findings showed conjugated hyperbilirubinaemia and significantly elevated transaminases (total bilirubin 83, conjugated 54, AST 993, ALT 714, GGT 155, AF 550). Complete blood count, coagulation profile, protein electrophoresis were initially normal. Viruses (HAV, HBV, HCV, HEV, CMV, EBV, Adenoviruses, Parvo B19) were excluded as a possible cause of hepatitis. In the next few days his clinical condition worsened, as well as his laboratory findings (total bilirubin 257, conjugated bilirubin 132, AST 2242, ALT 1399, GGT 91, AF 493, ammonia 119, significant deterioration of synthetic liver function). He developed hepatic encephalopathy and right-sided hemiconvulsions and was transferred to ICU where supportive treatment and plasmapheresis were started.

Liver biopsy showed extensive necrosis of parenchyma and moderate cholestasis. Three weeks after hospital admission, he underwent liver transplantation from cadaveric donor (II and III segment). During his first transplant, his native liver was removed completely, and he received cadaveric left lobe segment ll and lll).

Early postoperative period was complicated with development of portal vein thrombosis, so thrombectomy and portal vein neoanastomosis were performed on second posttransplantation day. During his stay in ICU he suffered gram positive sepsis and fungal pneumonia. A month after transplantation, he developed mechanical ileus due to numerous adhesions with bowel strangulation in the terminal ileum which were found on laparotomy. After two and half months, he was released from hospital. Extensive workup (screening for metabolic diseases, mitochondrial diseases autoimmune hepatitis, Wilson disease, drug intoxication) did not reveal the cause of acute liver failure in our patient.

In months following liver transplantation, he started to develop laboratory signs of chronic rejection, which was confirmed on liver biopsy so immunosupressive therapy was modified. He also started having focal seizures which were initially not considered epileptic since there was no EEG correlate, but during next few months he developed generalized symptomatic epilepsy and antiepileptic drugs were started. MRI showed diffuse cortical atrophy of the brain parenchyma, marked reduction in volume of white matter as well as gliosis in both frontal and temporoparietal lobes that could indicate the sequelae of acute hepatic encephalopathy, and 3.3 cm large arachnoid cyst in middle cranial fossa.

As epilepsy was refractory, several antiepileptic drugs were used to control it, and that was accompanied by further increase in liver enzymes. Second liver biopsy showed stationary finding mild chronic rejection. Calcineurin inhibitors which could hypothetically be epileptogenic were discontinued for two months, but it did not reduce seizure activity. Gradually, he started to develop signs of advanced liver disease (jaundice, itching) with further deterioration of liver function tests.

Two years after first liver transplant, he developed chronic rejection and was retransplanted (left cadaveric lobe from first transplant was removed and he received a new right cadaveric lobe). Surgery was successfully completed after 14 hours duration. Postoperative course was complicated with *Acinetobacter baumanii* sepsis, right sided pleural effusion and right lung atelectasis. Two weeks after re-transplantation, common bile duct stenosis was suspected based on ultrasound and MRCP findings, and was confirmed on relaparotomy. Resection of choledochojejunal anastomosis and formation of new terminolateral hepaticojejunal anastomosis was done. On 37th postoperative day he was transferred to ward, and the remaining course of stay proceed without further complications. Causes of graft failure were complex and included chronic rejection refractory to therapy, toxic effect of drugs and primary disease which was not well defined. Since he had several features (unusual facial characteristics, neurodevelopmental delay, autistic behavior, seizures and liver failure of undetermined cause) that could not fit in some well-known metabolic disease or syndrome, a whole genome sequencing was done.

A mutation of *ADNP* gene on the long arm of the 20th chromosome (20q13) corresponding to Helsmoortel-Van der AA syndrome (HVDAS) was discovered. In previously described patients with HVDAS syndrome there were no signs of liver disease. At the age of 4 years and 10 months, he was hospitalized in ICU due to SIRS and septic shock for 12 days. At the age of 5 years and 5 months, he had pneumonia caused by *Candida lusitanie* and prolonged enterocolitis. At 6 years of age, he was hospitalized due to elevation in transamninase (10x ULN) and GGT level (20xULN); coagulation profile and albumin levels were normal. He had reactivation of EBV infection (PCR EBV was positive in large number). CMV was negative.

At the age of 6 years and 2 months, another liver biopsy was done. PHD finding was inconclusive (received material was suboptimal, and could not confirm the diagnosis of EBV infection of the liver graft. Differential diagnosis considering morphological changes, could be an acute rejection -portal inflammation and damage to the bile ducts, but because of the disappearance of the bile ducts in certain areas could also be an early chronic rejection). Immunohistochemical analysis was negative for CMV. *In situ* hybridization EBV (EBER) was negative. 12 hours after liver biopsy he became febrile and developed septic shock and was transferred to ICU where he stayed until his death.

He was put on inotropic support and mechanical ventilation, broad spectrum antibiotics and antifungal therapy was started (*Candida parapsilosis* was isolated in repeated blood cultures), but he developed multiorgan failure and was put on hemodyalisis. His liver function also deteriorated. He had severe mucosal bleeding, received numerous blood transfusions, fresh frozen plasma, thrombocyte and fibrinogen transfusions; without improvements. His clinical condition was very difficult. He received 3 doses of Rituximab due to uncontrolled and progressive rise in EBV copies. Unfortunately, at the age of 6 years and 3 months, after one and half months stay in the ICU, he died.

## Legends to Supplementary Figures

**Supplementary Figure 1.**
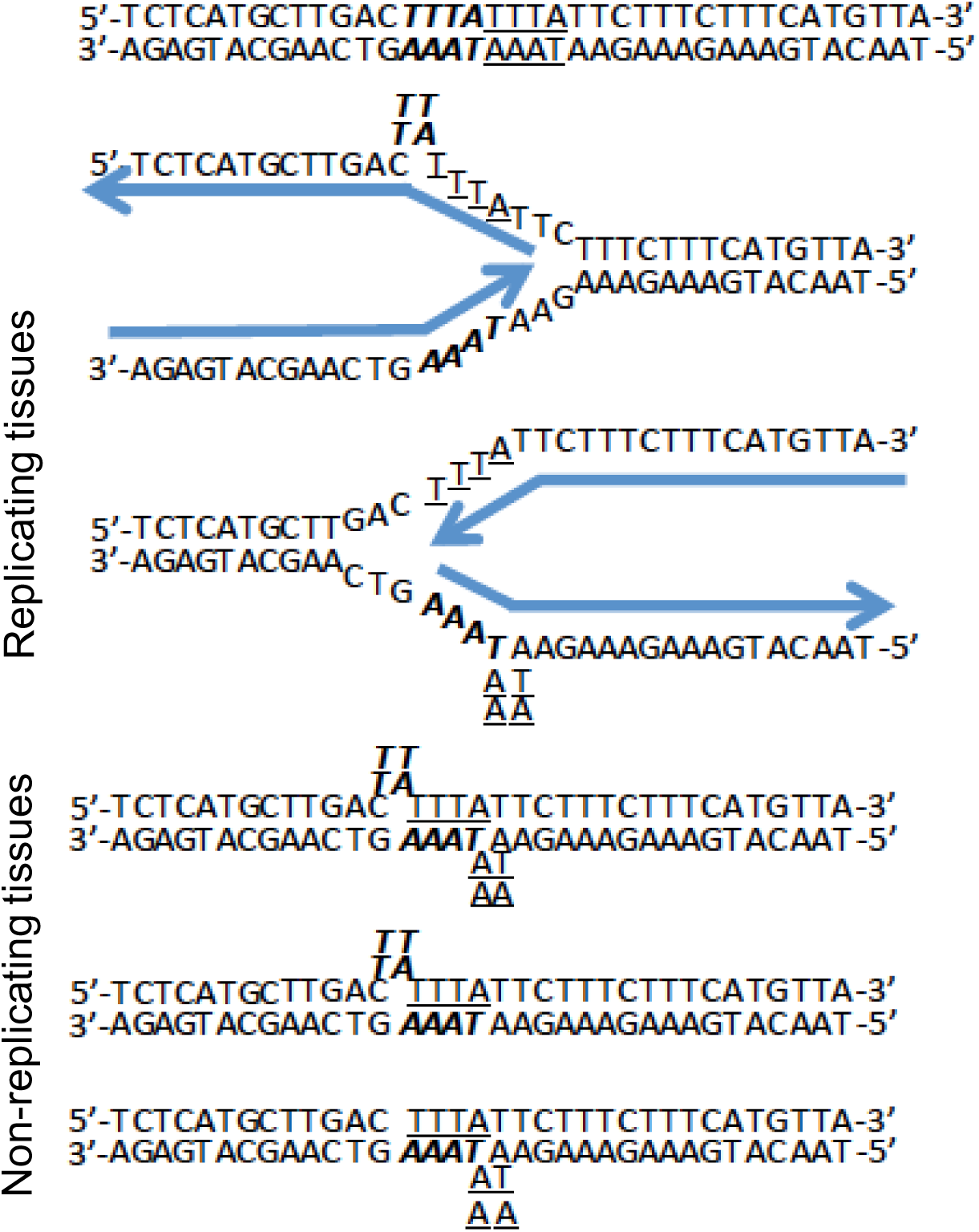
Stem-loop structure of the recurrent mutations during either DNA replication (proliferating tissues) or DNA repair (non-proliferating tissues, such as the CNS). The mutation hotspot is indicated in top: 5’ TTGAC**TTTA**TTTATTCTTTCTTTC-3’, where the bolded italicized **TTTA** sequences are the frequently deleted in patients.

**Supplementary Figure 2.**
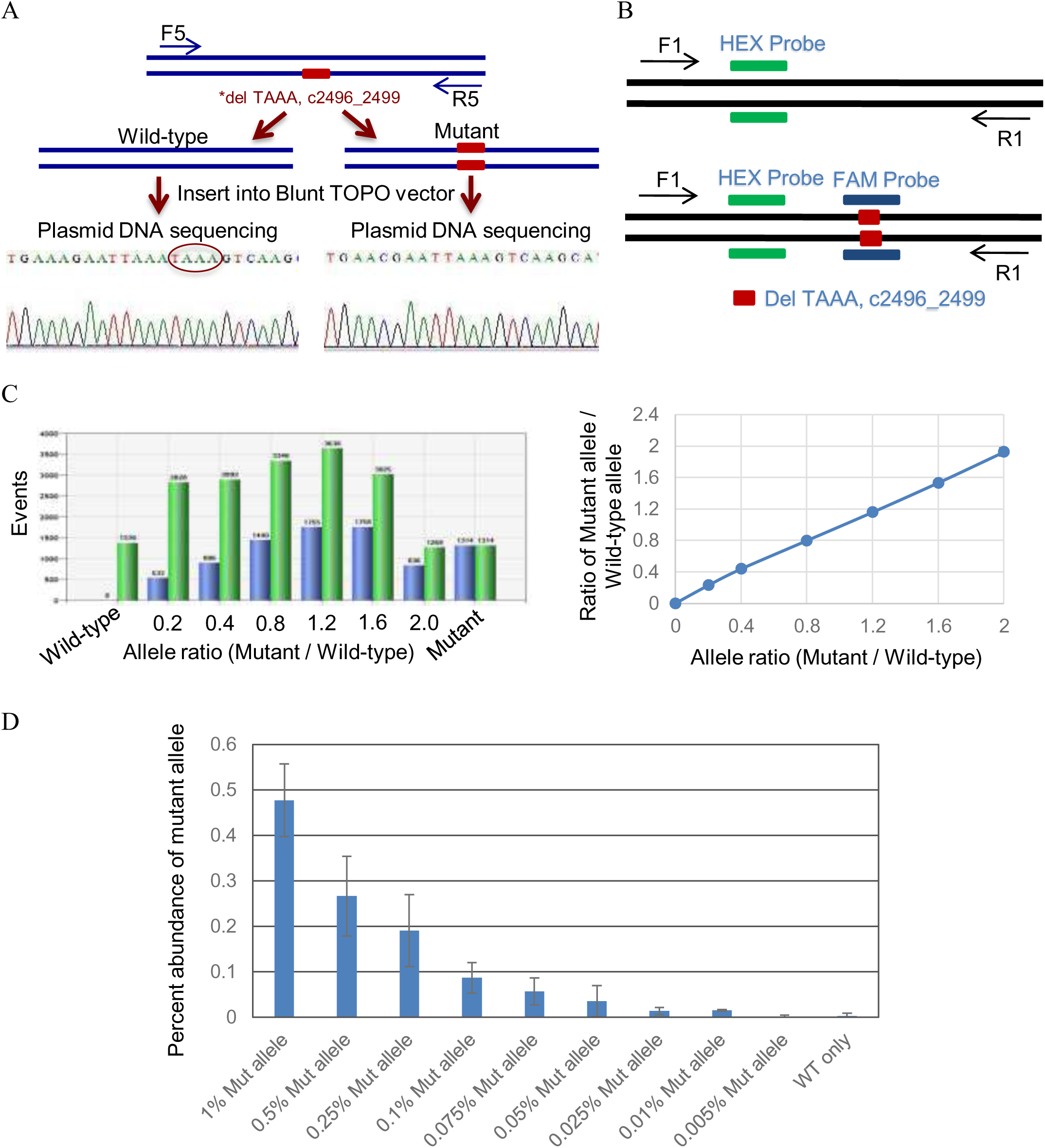
Digital droplet PCR (ddPCR) allows to quantify samples without using a standard curve. (A) Schematic illustration of generating homozygous *ADNP* mutant (c.2496-2499delTAAA) DNA containing Topo vector. (B) Schematic illustration for non-competing (hybrid) duplex reactions of ddPCR assay to generate standard curve. HEX probe (universal / reference probe) is designed upstream of mutation site (c.2496_2499TAAAdel), whereas FAM probe is specific for mutation region. Primer pair (F1/R1) amplifies a sequence covering both HEX and FAM probes specific sequences. (C) Histogram representing the number of events (positive droplets for FAM or HEX probes) (y-axis) in the indicated allelic ratio (x-axis) (left panel). Right panel shows standard curve (linear regression), which is drawn based upon mixed ratios of DNAs (*x-axis*) and ratio of FAM and HEX positive droplets (*y-axis*). (D) Sensitive detection of mutant alleles. Droplet digital PCR assay against c.2496-2499delTAAA *ADNP* mutation can detect the mutant allele down to a frequency of 0.01% (samples run in at least triplicate). The background positive level ranges from 0.0024% to 0, allowing for sensitive detection of the mutant allele at 0.01%.

**Supplementary Figure 3.**
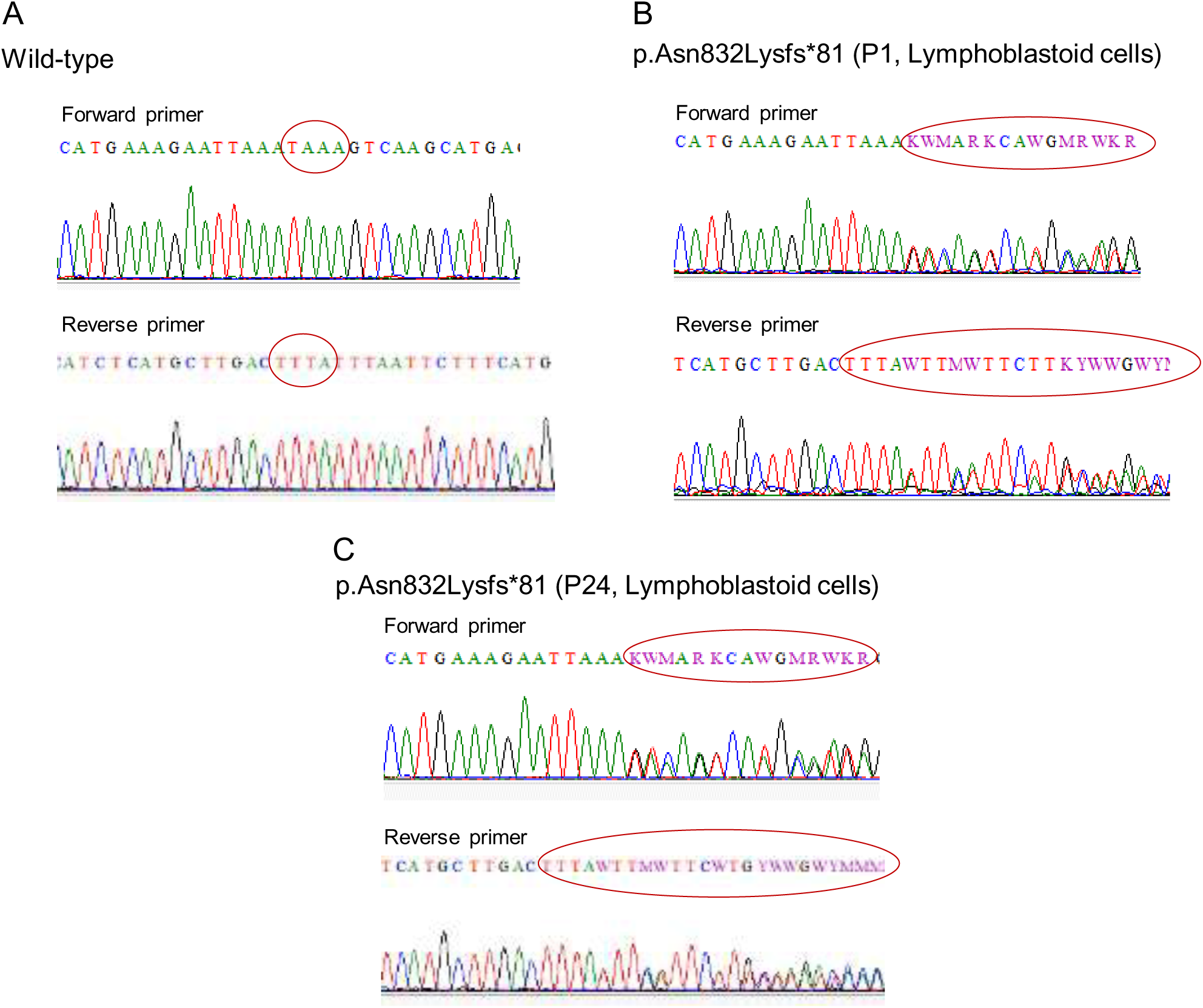
Validation for c.2496_2499TAAAdel mutation. *De novo* 4 bp *ADNP* frameshift deletion (c.2496_2499TAAAdel) detected in patient 1 was confirmed by Sanger sequencing directly from genomic DNA. Sequence chromatographs covering the *ADNP* codon in the non-patient wild-type DNA (A), DNA isolated from lymphoblastoid cell line derived from patient 1 (B) and patient 24 (C).

**Supplementary Figure 4.**
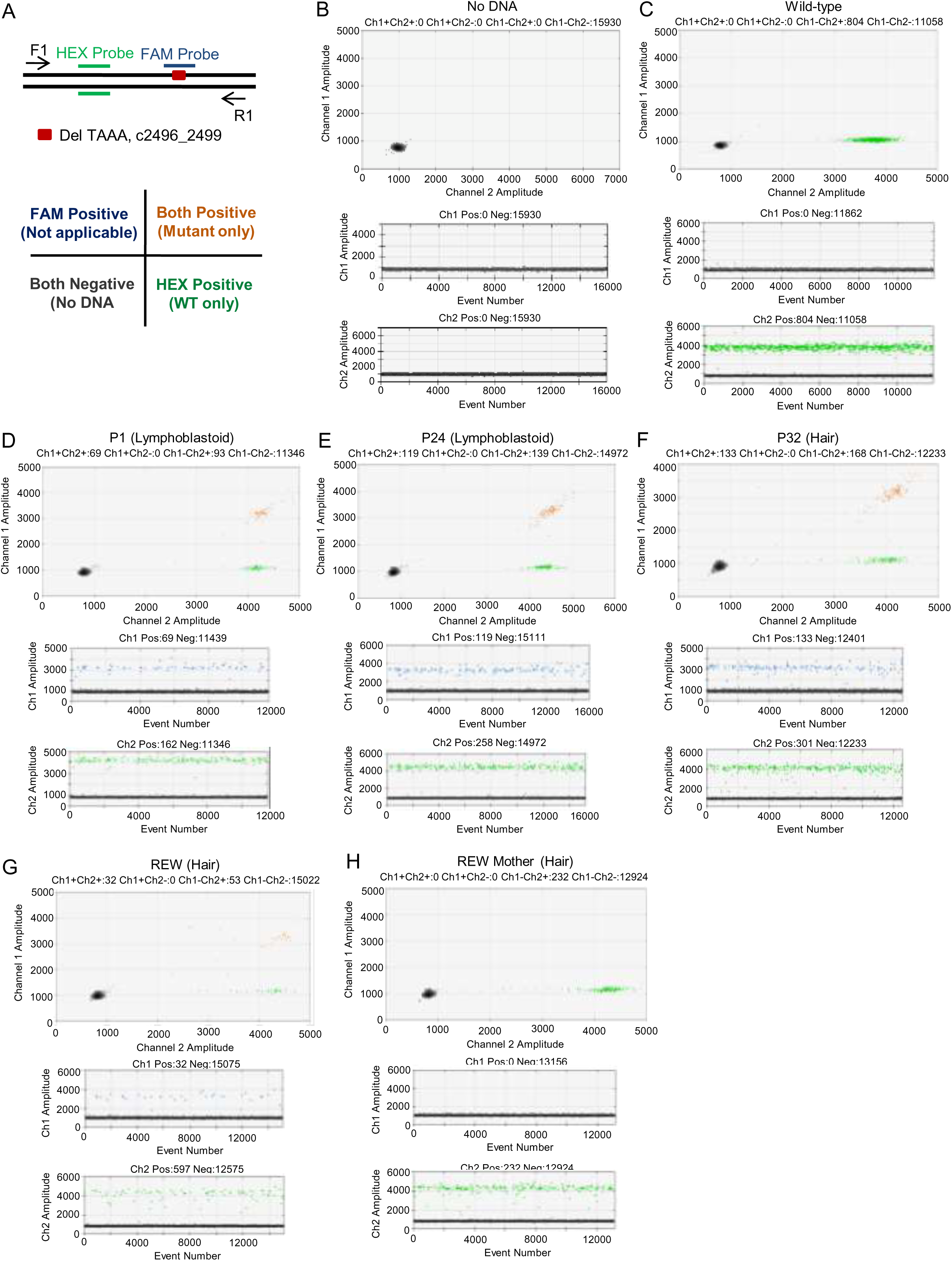
Molecular evaluation of *ADNP* mutation (c.2496_2499TAAAdel) in different neural tissues, lymphoblastoid cells and iPSCs clones. (A) Schematic illustration for non-competing (hybrid) duplex reactions of ddPCR assay to evaluate mutation load in different tissues, and lymphoblastoid cells (left panel). HEX probe (universal / reference probe) is designed upstream of mutation site (c.2496_2499TAAAdel), whereas FAM probe is specific for mutation region. Primer pair (F1/R1) amplifies a sequence covering both HEX and FAM probes specific sequences. Only three clusters are visible in the non-competing (hybrid) duplex reactions (right panel). (B-F) Upper panel: 2-D plot showing amplification of *ADNP* gene in the Blank (B), Wild-type (C), P1-Lymphoblastoid cells (D), P24- Lymphoblastoid cells (E), P32-hair (F), P100-hair (G) and P100-Mother-hair (H) DNAs. Lower Panel: 1-D plot showing *ADNP* gene amplification. A well-defined separation between negative droplets and positive droplets was seen.

**Supplementary Figure 5.**
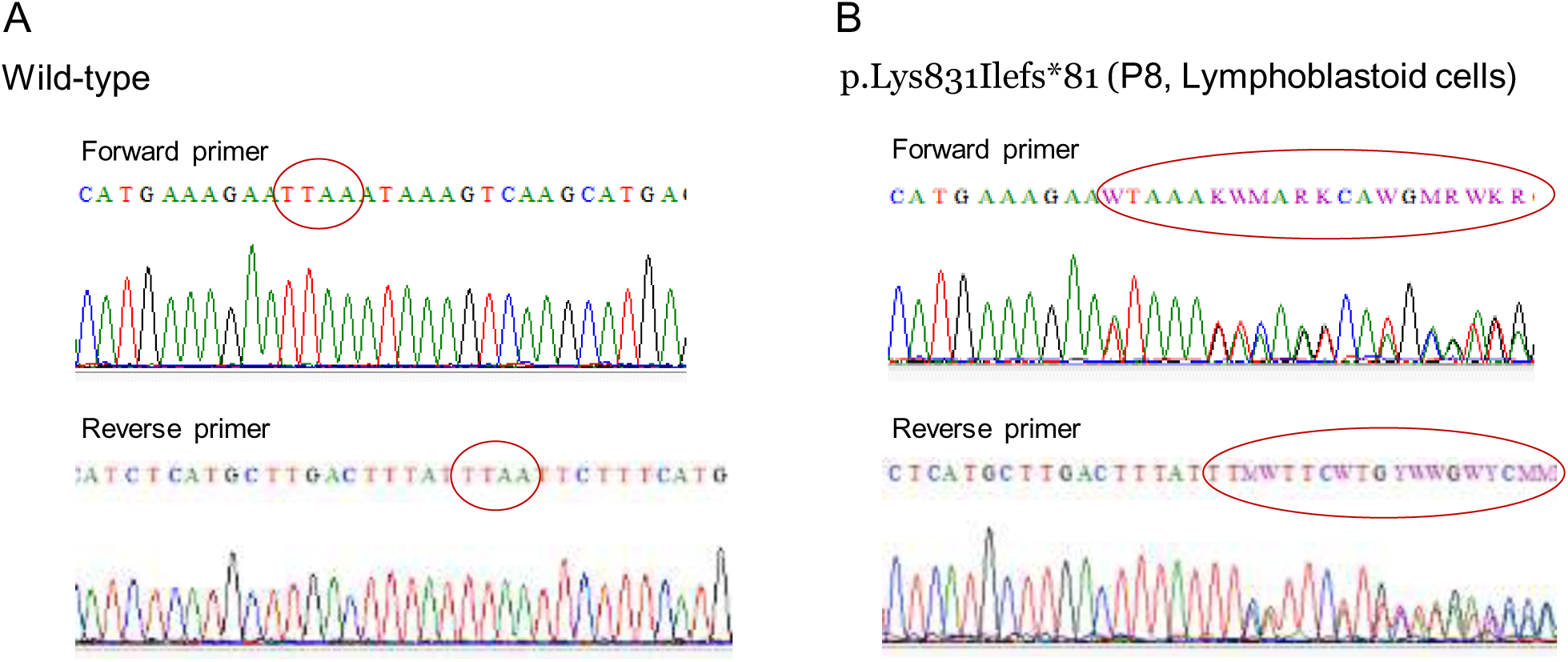
Validation for c.2491_2494TTAAdel mutation. *De novo* 4 bp *ADNP* frameshift deletion (c.2491_2494TTAAdel) detected in patient 8 was confirmed by Sanger sequencing directly from genomic DNA. Sequence chromatograph covering the *ADNP* codon in the non-patient wild-type DNA (A) and DNA isolated from lymphoblastoid cell line derived from patient 8 (B).

**Supplementary Figure 6.**
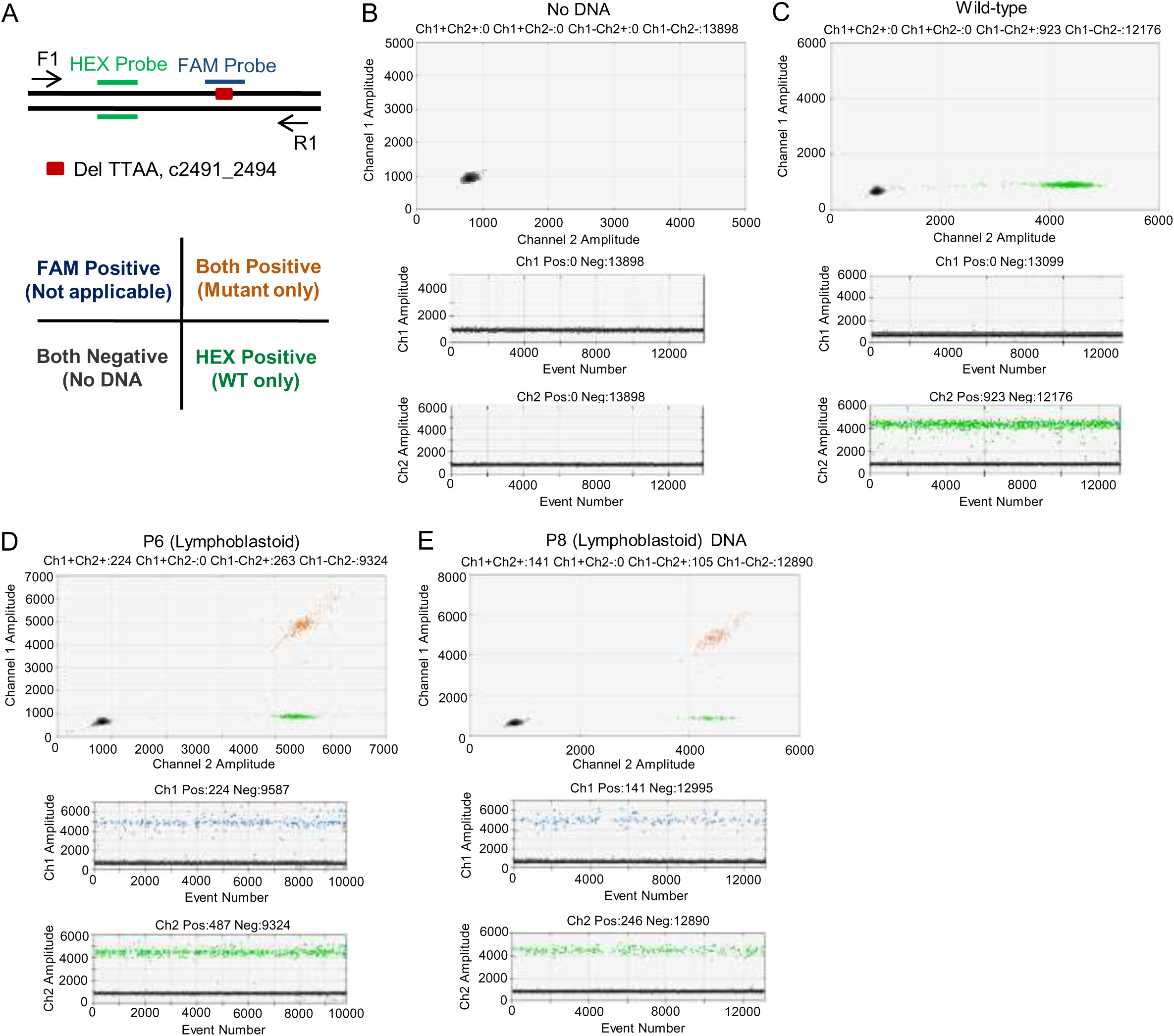
Degree of mosaicism and molecular evaluation of *ADNP* mutation (c.2491_2494TTAAdel) in lymphoblastoid cells. (A) Schematic illustration for non-competing (hybrid) duplex reactions of ddPCR assay to evaluate mutation load in lymphoblastoid cells and iPSC clones (left panel). HEX probe (universal / reference probe) is designed upstream of mutation site (c.2491_2494TTAAdel), whereas FAM probe is specific for mutation region. Primer pair (F1/R1) amplifies a sequence covering both HEX and FAM probes specific sequences. Only three clusters are visible in the non-competing (hybrid) duplex reactions (right panel). (B-E) Upper panel: 2-D plot showing amplification of *ADNP* gene in the Blank (B), Wild-type (C), P6-Lymphoblastoid (D), and P8-Lymphoblastoid cells (E) DNAs. Lower Panel: 1-D plot showing *ADNP* gene amplification. A well-defined separation between negative droplets and positive droplets for both FAM and HEX probes was seen.

**Supplementary Figure 7.**
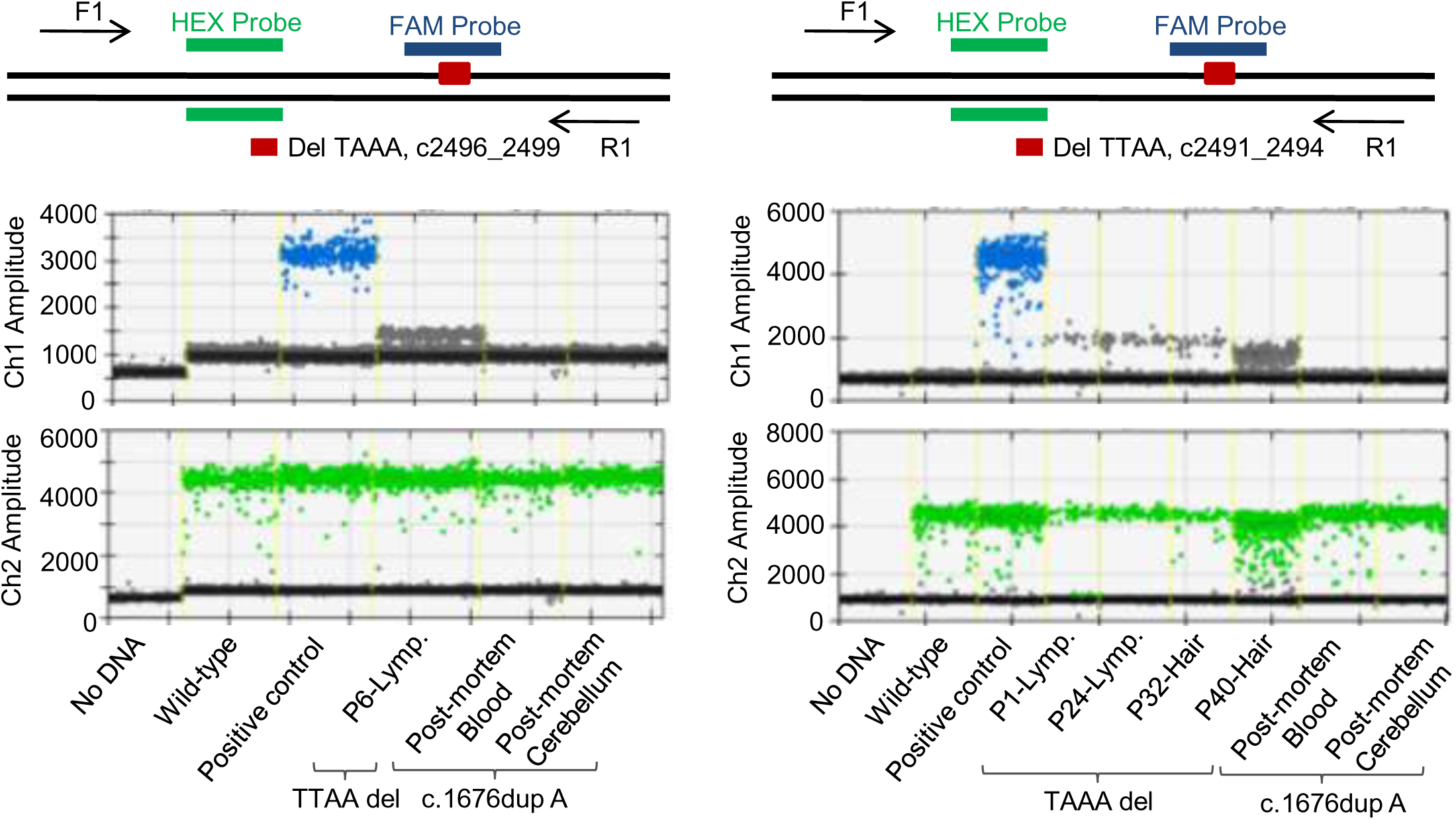
*ADNP* patient tissues or patient-derived lymphoblastoid cells are cross-contamination free. (A) Upper panel: Schematic illustration for non-competing (hybrid) duplex reactions of ddPCR assay to evaluate (c.2496_2499TAAAdel (B) and c.2491_2494TTAAdel (B) mutation loads in different tissues, and lymphoblastoid cells. Lower panel: 1D plots for FAM (blue) and HEX (green) positive droplets in the indicative cells and tissues.

**Supplementary Figure 8.**
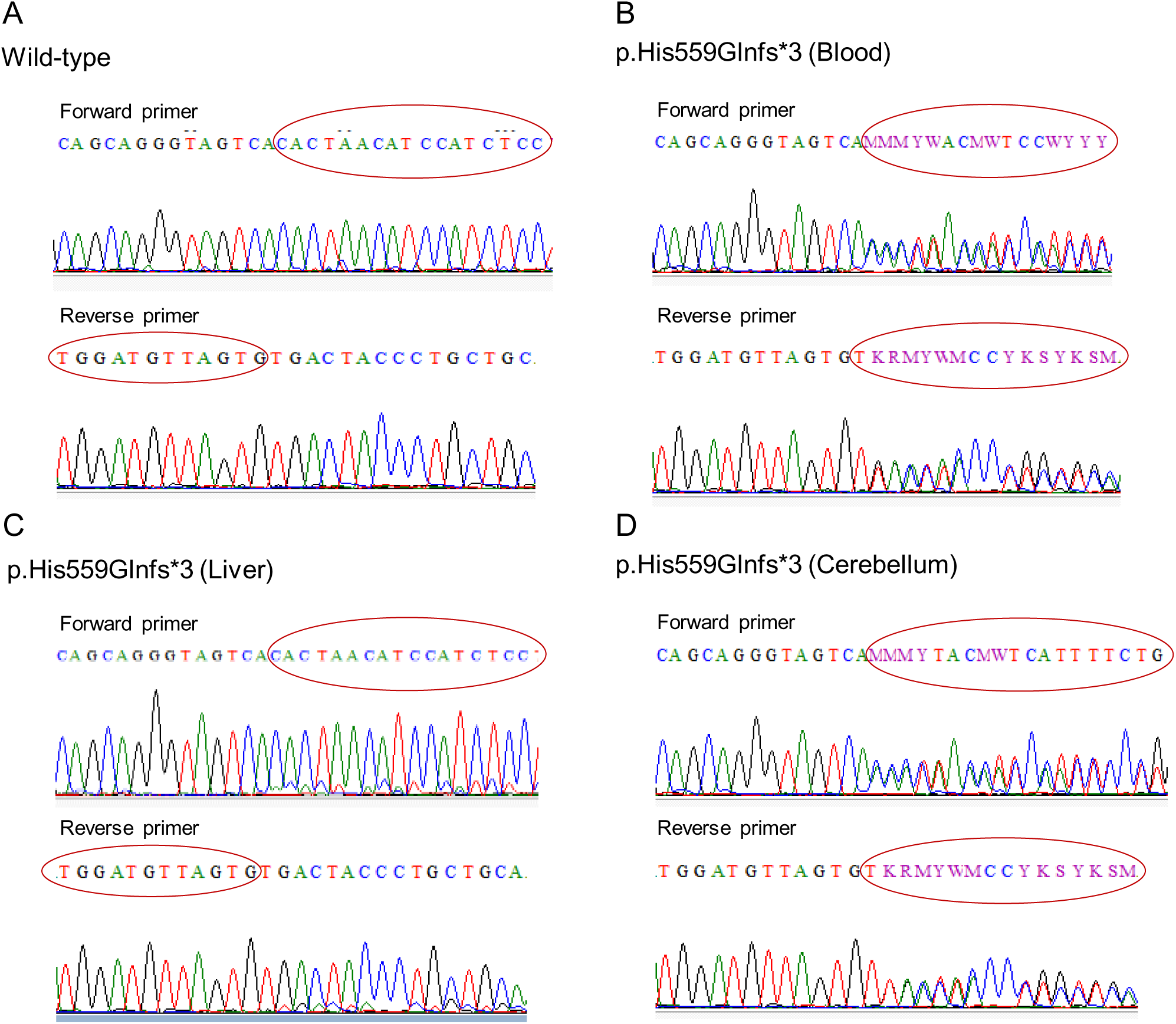
Validation for c.1676dupA mutation. *De novo* frameshift mutation in the *ADNP* gene at His599 (c.1676dup) in the post-mortem tissues obtained from a deceased *ADNP* patient were confirmed by Sanger sequencing directly from genomic DNA. Sequence chromatograph covering the *ADNP* codon in the non-patient wild-type DNA (A) and DNA isolated from Blood (B), Liver (C) and Cerebellum (D) from a deceased *ADNP* patient.

**Supplementary Figure 9.**
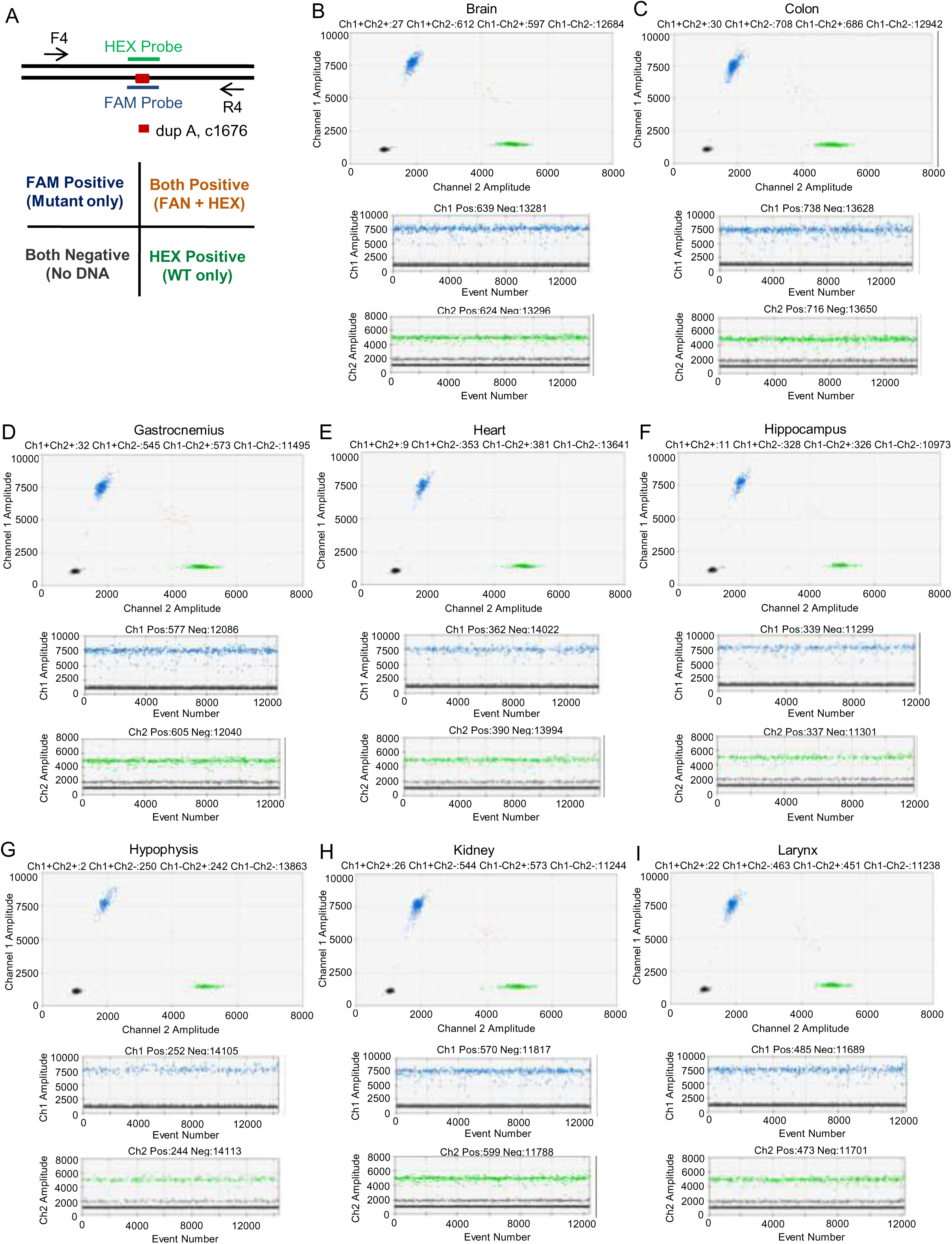
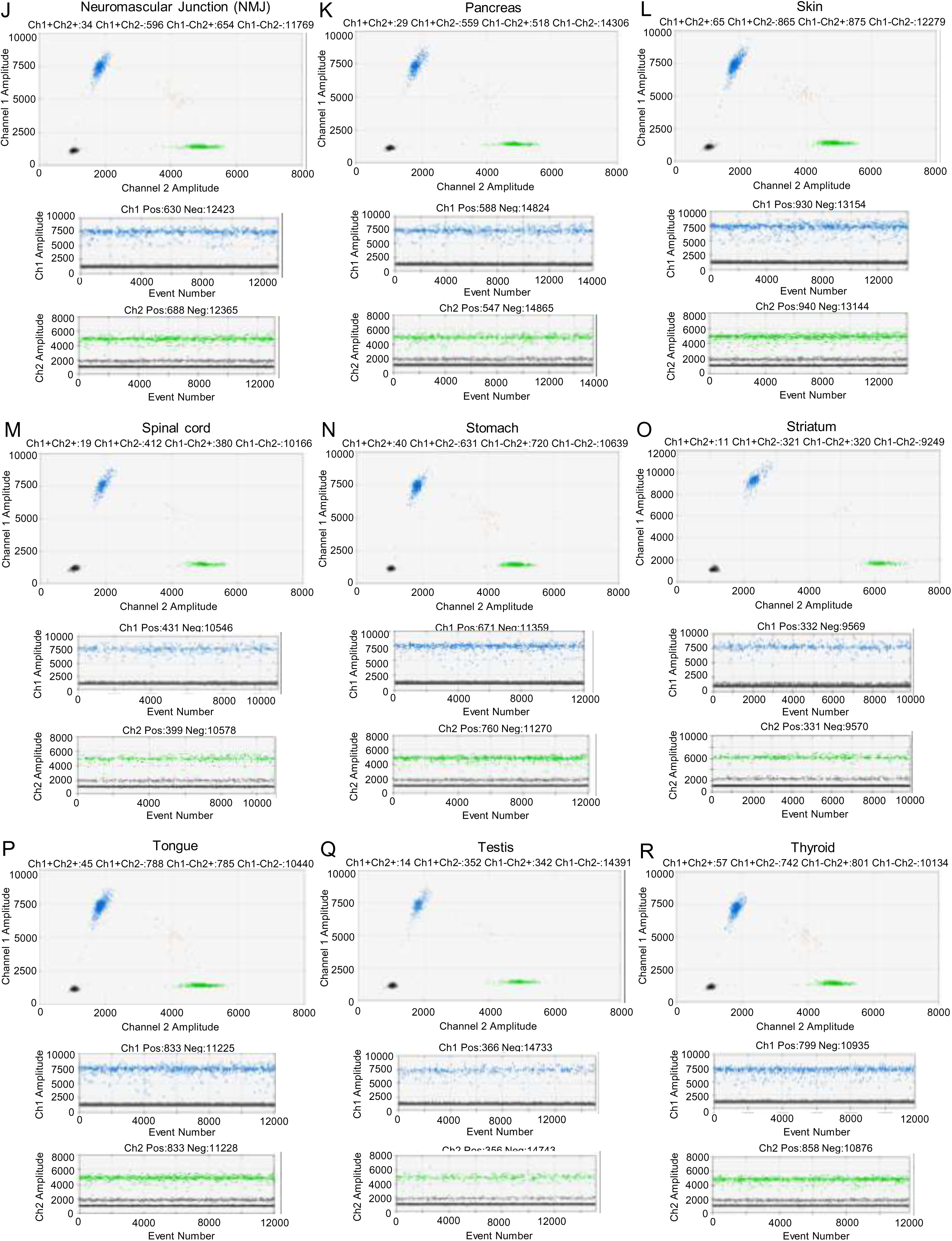
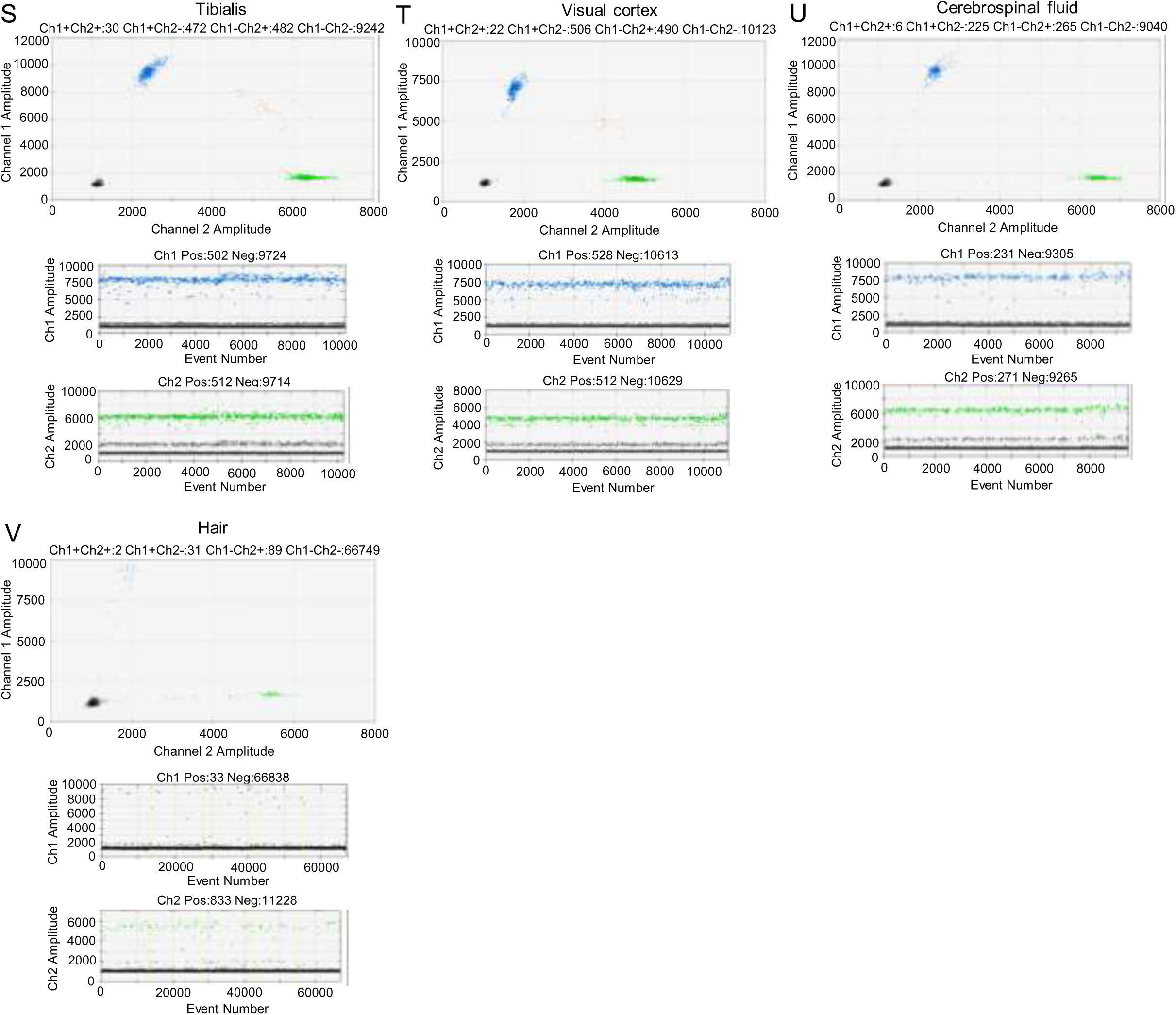
Molecular evaluation of *ADNP* mutation (c.1676dupA) in the post-mortem tissues obtained from a deceased *ADNP* patient. (A) Schematic illustration for competing duplex reactions of ddPCR assay to evaluate mutation load in the post-mortem tissues obtained from a deceased *ADNP* patient (upper panel). HEX probe (universal / reference probe) is designed for the abundant or wild-type sequence, whereas FAM probe is designed for the mutant sequence (c.1676dupA). Primer pair (F4/R4) amplifies a sequence covering c.1676 of *ADNP* gene. Unlike non-competing (hybrid) duplex reactions, four clusters are visible in the competing duplex reactions (lower panel). For the analysis of a probe-competing duplex reactions to quantify a single base substitution, cross-hybridization of the probes caused by either mismatched of the probes or filter bleed-though can be visualized as a ‘leaning’ (blue) or ‘lifting’ (green) of the single positive clusters. (B-V) Upper panel: 2-D plot showing amplification of *ADNP* gene in the Brain (B), Colon (C), Gastrocnemius (D), Heart (E), Hippocampus (F), Hypophysis (G), Kidney (H), Larynx (I), NMJ (J), Pancreas (K), Skin (L), Spinal cord (M), Stomach (N), Striatum (O), Tongue (P), Testis (Q), Thyroid (R), Tibialis (S), Visual cortex (T), Cerebrospinal fluid (U) and Hair (V) DNAs. Lower Panel: 1-D plot showing *ADNP* gene amplification. A well-defined separation between negative droplets and positive droplets was seen.

**Supplementary Table 1.**
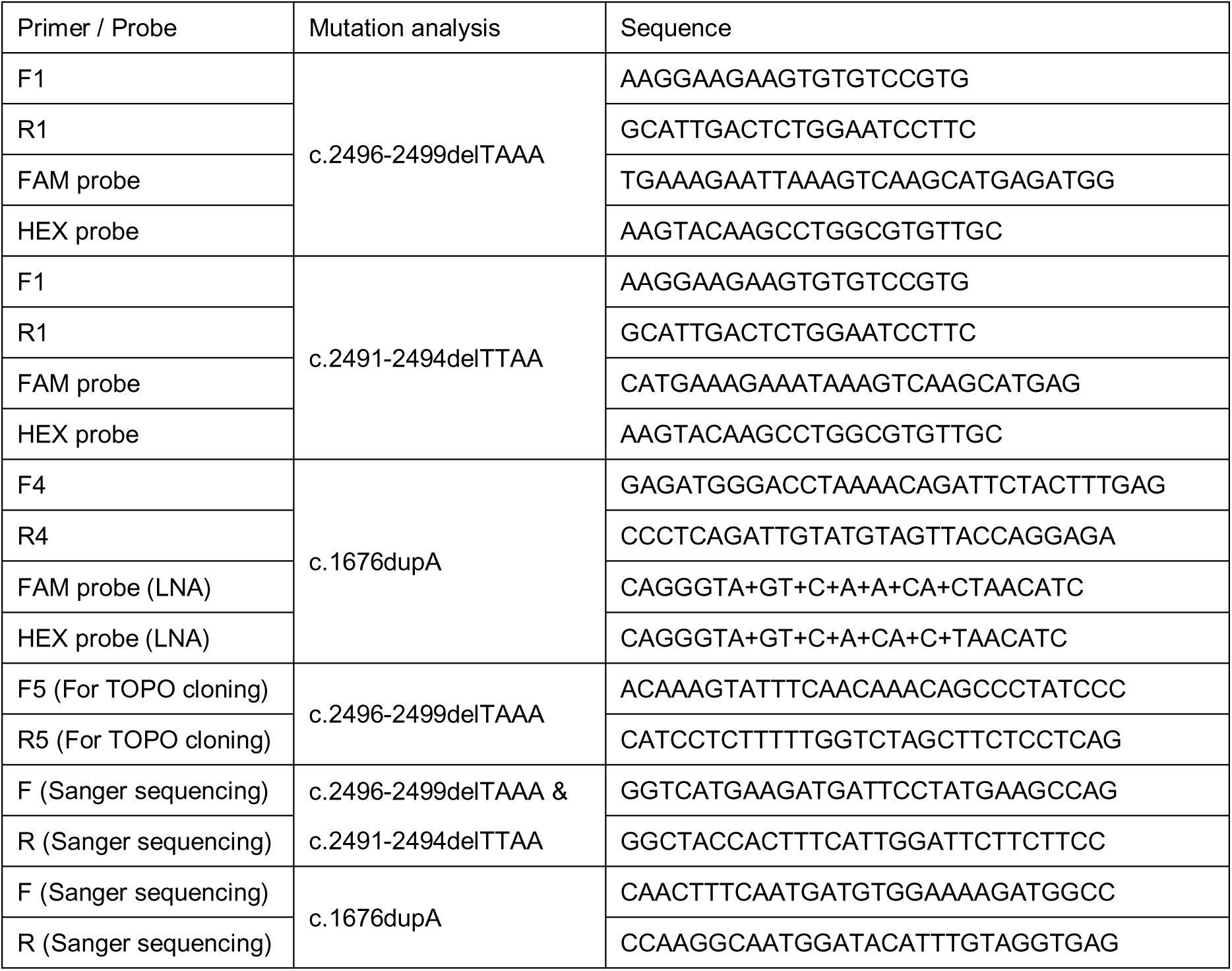
List of primers and probes used in this study.

**Supplementary Table 2.** Forensic analysis of post-mortem tissues used in this study.

| Sample Name | Panel | Marker | Allele 1 | Allele 2 | Allele 3 | Allele 4 |
| --- | --- | --- | --- | --- | --- | --- |
| 01-Blood | GenePrint | AMEL | X | Y | - | - |
| 02-Brain Stem | GenePrint | AMEL | X | Y | - | - |
| 03-Cerebellum | GenePrint | AMEL | X | Y | - | - |
| 04-Colon | GenePrint | AMEL | X | Y | - | - |
| 05-CSF | GenePrint | AMEL | X | Y | - | - |
| 06-Gastrocnemius | GenePrint | AMEL | X | Y | - | - |
| 07-Heart | GenePrint | AMEL | X | Y | - | - |
| 08-Hippocampus | GenePrint | AMEL | X | Y | - | - |
| 09-Hypophysis | GenePrint | AMEL | X | Y | - | - |
| 10-Kidney | GenePrint | AMEL | X | Y | - | - |
| 11-Larynx | GenePrint | AMEL | X | Y | - | - |
| <u>12-Liver</u> | <u>GenePrint</u> | <u>AMEL</u> | <u>X</u> | <u>Y</u> | - | - |
| <u>13-NMJ</u> | <u>GenePrint</u> | <u>AMEL</u> | <u>X</u> | <u>Y</u> | - | - |
| <u>14-Pancreas</u> | <u>GenePrint</u> | <u>AMEL</u> | <u>X</u> | <u>Y</u> | - | - |
| <u>15-Skin</u> | <u>GenePrint</u> | <u>AMEL</u> | <u>X</u> | <u>Y</u> | - | - |
| <u>16-Spinal_Cord</u> | <u>GenePrint</u> | <u>AMEL</u> | <u>X</u> | <u>Y</u> | - | - |
| <u>17-Stomach</u> | <u>GenePrint</u> | <u>AMEL</u> | <u>X</u> | <u>Y</u> | - | - |
| <u>18-Striatum</u> | <u>GenePrint</u> | <u>AMEL</u> | <u>X</u> | <u>Y</u> | - | - |
| <u>19-Tongue</u> | <u>GenePrint</u> | <u>AMEL</u> | <u>X</u> | <u>Y</u> | - | - |
| <u>20-Testis</u> | <u>GenePrint</u> | <u>AMEL</u> | <u>X</u> | <u>Y</u> | - | - |
| <u>21-Thyroid</u> | <u>GenePrint</u> | <u>AMEL</u> | <u>X</u> | <u>Y</u> | - | - |
| <u>22-Tibialis</u> | <u>GenePrint</u> | <u>AMEL</u> | <u>X</u> | <u>Y</u> | - | - |
| <u>23-Visual_Cortex</u> | <u>GenePrint</u> | <u>AMEL</u> | <u>X</u> | <u>Y</u> | - | - |
| <u>01-Blood</u> | <u>GenePrint</u> | <u>CSF1PO</u> | <u>9</u> | <u>12</u> | - | - |
| <u>02-Brain_Stem</u> | <u>GenePrint</u> | <u>CSF1PO</u> | <u>9</u> | <u>12</u> | - | - |
| <u>03-Cerebellum</u> | <u>GenePrint</u> | <u>CSF1PO</u> | <u>9</u> | <u>12</u> | - | - |
| <u>04-Colon</u> | <u>GenePrint</u> | <u>CSF1PO</u> | <u>9</u> | <u>12</u> | - | - |
| <u>05-CSF</u> | <u>GenePrint</u> | <u>CSF1PO</u> | <u>9</u> | <u>12</u> | - | - |
| <u>06-Gastrocnemius</u> | <u>GenePrint</u> | <u>CSF1PO</u> | <u>9</u> | <u>12</u> | - | - |
| <u>07-Heart</u> | <u>GenePrint</u> | <u>CSF1PO</u> | <u>9</u> | <u>12</u> | - | - |
| <u>08-Hippocampus</u> | <u>GenePrint</u> | <u>CSF1PO</u> | <u>9</u> | <u>12</u> | - | - |
| <u>09-Hypophysis</u> | <u>GenePrint</u> | <u>CSF1PO</u> | <u>9</u> | <u>12</u> | - | - |
| <u>10-Kidney</u> | <u>GenePrint</u> | <u>CSF1PO</u> | <u>9</u> | <u>12</u> | - | - |
| <u>11-Larynx</u> | <u>GenePrint</u> | <u>CSF1PO</u> | <u>9</u> | <u>12</u> | - | - |
| <u>12-Liver</u> | <u>GenePrint</u> | <u>CSF1PO</u> | <u>9</u> | <u>10</u> | <u>12</u> | - |
| <u>13-NMJ</u> | <u>GenePrint</u> | <u>CSF1PO</u> | <u>9</u> | <u>12</u> | - | - |
| <u>14-Pancreas</u> | <u>GenePrint</u> | <u>CSF1PO</u> | <u>9</u> | <u>12</u> | - | - |
| <u>15-Skin</u> | <u>GenePrint</u> | <u>CSF1PO</u> | <u>9</u> | <u>12</u> | - | - |
| <u>16-Spinal_Cord</u> | <u>GenePrint</u> | <u>CSF1PO</u> | <u>9</u> | <u>12</u> | - | - |
| <u>17-Stomach</u> | <u>GenePrint</u> | <u>CSF1PO</u> | <u>9</u> | <u>12</u> | - | - |
| <u>18-Striatum</u> | <u>GenePrint</u> | <u>CSF1PO</u> | <u>9</u> | <u>12</u> | - | - |
| <u>19-Tongue</u> | <u>GenePrint</u> | <u>CSF1PO</u> | <u>9</u> | <u>12</u> | - | - |
| <u>20-Testis</u> | <u>GenePrint</u> | <u>CSF1PO</u> | <u>9</u> | <u>12</u> | - | - |
| <u>21-Thyroid</u> | <u>GenePrint</u> | <u>CSF1PO</u> | <u>9</u> | <u>12</u> | - | - |
| <u>22-Tibialis</u> | <u>GenePrint</u> | <u>CSF1PO</u> | <u>9</u> | <u>12</u> | - | - |
| <u>23-Visual_Cortex</u> | <u>GenePrint</u> | <u>CSF1PO</u> | <u>9</u> | <u>12</u> | - | - |
| <u>01-Blood</u> | <u>GenePrint</u> | <u>D13S317</u> | <u>8</u> | <u>9</u> | - | - |
| <u>02-Brain_Stem</u> | <u>GenePrint</u> | <u>D13S317</u> | <u>8</u> | <u>9</u> | - | - |
| <u>03-Cerebellum</u> | <u>GenePrint</u> | <u>D13S317</u> | <u>8</u> | <u>9</u> | - | - |
| <u>04-Colon</u> | <u>GenePrint</u> | <u>D13S317</u> | <u>8</u> | <u>9</u> | - | - |
| <u>05-CSF</u> | <u>GenePrint</u> | <u>D13S317</u> | <u>8</u> | <u>9</u> | - | - |
| <u>06-Gastrocnemius</u> | <u>GenePrint</u> | <u>D13S317</u> | <u>8</u> | <u>9</u> | - | - |
| <u>07-Heart</u> | <u>GenePrint</u> | <u>D13S317</u> | <u>8</u> | <u>9</u> | - | - |
| <u>08-Hippocampus</u> | <u>GenePrint</u> | <u>D13S317</u> | <u>8</u> | <u>9</u> | - | - |
| <u>09-Hypophysis</u> | <u>GenePrint</u> | <u>D13S317</u> | <u>8</u> | <u>9</u> | - | - |
| <u>10-Kidney</u> | <u>GenePrint</u> | <u>D13S317</u> | <u>8</u> | <u>9</u> | - | - |
| <u>11-Larynx</u> | <u>GenePrint</u> | <u>D13S317</u> | <u>8</u> | <u>9</u> | - | - |
| <u>12-Liver</u> | <u>GenePrint</u> | <u>D13S317</u> | <u>8</u> | <u>9</u> | <u>13</u> | <u>14</u> |
| <u>13-NMJ</u> | <u>GenePrint</u> | <u>D13S317</u> | <u>8</u> | <u>9</u> | - | - |
| <u>14-Pancreas</u> | <u>GenePrint</u> | <u>D13S317</u> | <u>8</u> | <u>9</u> | - | - |
| <u>15-Skin</u> | <u>GenePrint</u> | <u>D13S317</u> | <u>8</u> | <u>9</u> | - | - |
| <u>16-Spinal_Cord</u> | <u>GenePrint</u> | <u>D13S317</u> | <u>8</u> | <u>9</u> | - | - |
| <u>17-Stomach</u> | <u>GenePrint</u> | <u>D13S317</u> | <u>8</u> | <u>9</u> | - | - |
| <u>18-Striatum</u> | <u>GenePrint</u> | <u>D13S317</u> | <u>8</u> | <u>9</u> | - | - |
| <u>19-Tongue</u> | <u>GenePrint</u> | <u>D13S317</u> | <u>8</u> | <u>9</u> | - | - |
| <u>20-Testis</u> | <u>GenePrint</u> | <u>D13S317</u> | <u>8</u> | <u>9</u> | - | - |
| <u>21-Thyroid</u> | <u>GenePrint</u> | <u>D13S317</u> | <u>8</u> | <u>9</u> | - | - |
| <u>22-Tibialis</u> | <u>GenePrint</u> | <u>D13S317</u> | <u>8</u> | <u>9</u> | - | - |
| <u>23-Visual_Cortex</u> | <u>GenePrint</u> | <u>D13S317</u> | <u>8</u> | <u>9</u> | - | - |
| <u>01-Blood</u> | <u>GenePrint</u> | <u>D16S539</u> | <u>9</u> | <u>12</u> | - | - |
| <u>02-Brain_Stem</u> | <u>GenePrint</u> | <u>D16S539</u> | <u>9</u> | <u>12</u> | - | - |
| <u>03-Cerebellum</u> | <u>GenePrint</u> | <u>D16S539</u> | <u>9</u> | <u>12</u> | - | - |
| <u>04-Colon</u> | <u>GenePrint</u> | <u>D16S539</u> | <u>9</u> | <u>12</u> | - | - |
| <u>05-CSF</u> | <u>GenePrint</u> | <u>D16S539</u> | <u>9</u> | <u>12</u> | - | - |
| <u>06-Gastrocnemius</u> | <u>GenePrint</u> | <u>D16S539</u> | <u>9</u> | <u>12</u> | - | - |
| <u>07-Heart</u> | <u>GenePrint</u> | <u>D16S539</u> | <u>9</u> | <u>12</u> | - | - |
| <u>08-Hippocampus</u> | <u>GenePrint</u> | <u>D16S539</u> | <u>9</u> | <u>12</u> | - | - |
| <u>09-Hypophysis</u> | <u>GenePrint</u> | <u>D16S539</u> | <u>9</u> | <u>12</u> | - | - |
| <u>10-Kidney</u> | <u>GenePrint</u> | <u>D16S539</u> | <u>9</u> | <u>12</u> | - | - |
| <u>11-Larynx</u> | <u>GenePrint</u> | <u>D16S539</u> | <u>9</u> | <u>12</u> | - | - |
| <u>12-Liver</u> | <u>GenePrint</u> | <u>D16S539</u> | <u>8</u> | <u>9</u> | <u>10</u> | <u>12</u> |
| <u>13-NMJ</u> | <u>GenePrint</u> | <u>D16S539</u> | <u>9</u> | <u>12</u> | - | - |
| <u>14-Pancreas</u> | <u>GenePrint</u> | <u>D16S539</u> | <u>9</u> | <u>12</u> | - | - |
| <u>15-Skin</u> | <u>GenePrint</u> | <u>D16S539</u> | <u>9</u> | <u>12</u> | - | - |
| <u>16-Spinal_Cord</u> | <u>GenePrint</u> | <u>D16S539</u> | <u>9</u> | <u>12</u> | - | - |
| <u>17-Stomach</u> | <u>GenePrint</u> | <u>D16S539</u> | <u>9</u> | <u>12</u> | - | - |
| <u>18-Striatum</u> | <u>GenePrint</u> | <u>D16S539</u> | <u>9</u> | <u>12</u> | - | - |
| <u>19-Tongue</u> | <u>GenePrint</u> | <u>D16S539</u> | <u>9</u> | <u>12</u> | - | - |
| <u>20-Testis</u> | <u>GenePrint</u> | <u>D16S539</u> | <u>9</u> | <u>12</u> | - | - |
| <u>21-Thyroid</u> | <u>GenePrint</u> | <u>D16S539</u> | <u>9</u> | <u>12</u> | - | - |
| <u>22-Tibialis</u> | <u>GenePrint</u> | <u>D16S539</u> | <u>9</u> | <u>12</u> | - | - |
| <u>23-Visual Cortex</u> | <u>GenePrint</u> | <u>D16S539</u> | <u>9</u> | <u>12</u> | - | - |
| <u>01-Blood</u> | <u>GenePrint</u> | <u>D21S11</u> | <u>30.2</u> | <u>31.2</u> | - | - |
| <u>02-Brain Stem</u> | <u>GenePrint</u> | <u>D21S11</u> | <u>30.2</u> | <u>31.2</u> | - | - |
| <u>03-Cerebellum</u> | <u>GenePrint</u> | <u>D21S11</u> | <u>30.2</u> | <u>31.2</u> | - | - |
| <u>04-Colon</u> | <u>GenePrint</u> | <u>D21S11</u> | <u>30.2</u> | <u>31.2</u> | - | - |
| <u>05-CSF</u> | <u>GenePrint</u> | <u>D21S11</u> | <u>30.2</u> | <u>31.2</u> | - | - |
| <u>06-Gastrocnemius</u> | <u>GenePrint</u> | <u>D21S11</u> | <u>30.2</u> | <u>31.2</u> | - | - |
| <u>07-Heart</u> | <u>GenePrint</u> | <u>D21S11</u> | <u>30.2</u> | <u>31.2</u> | - | - |
| <u>08-Hippocampus</u> | <u>GenePrint</u> | <u>D21S11</u> | <u>30.2</u> | <u>31.2</u> | - | - |
| <u>09-Hypophysis</u> | <u>GenePrint</u> | <u>D21S11</u> | <u>30.2</u> | <u>31.2</u> | - | - |
| <u>10-Kidney</u> | <u>GenePrint</u> | <u>D21S11</u> | <u>30.2</u> | <u>31.2</u> | - | - |
| <u>11-Larynx</u> | <u>GenePrint</u> | <u>D21S11</u> | <u>30.2</u> | <u>31.2</u> | - | - |
| <u>12-Liver</u> | <u>GenePrint</u> | <u>D21S11</u> | <u>28</u> | <u>30.2</u> | <u>31.2</u> | - |
| <u>13-NMJ</u> | <u>GenePrint</u> | <u>D21S11</u> | <u>30.2</u> | <u>31.2</u> | - | - |
| <u>14-Pancreas</u> | <u>GenePrint</u> | <u>D21S11</u> | <u>30.2</u> | <u>31.2</u> | - | - |
| <u>15-Skin</u> | <u>GenePrint</u> | <u>D21S11</u> | <u>30.2</u> | <u>31.2</u> | - | - |
| <u>16-Spinal Cord</u> | <u>GenePrint</u> | <u>D21S11</u> | <u>30.2</u> | <u>31.2</u> | - | - |
| <u>17-Stomach</u> | <u>GenePrint</u> | <u>D21S11</u> | <u>30.2</u> | <u>31.2</u> | - | - |
| <u>18-Striatum</u> | <u>GenePrint</u> | <u>D21S11</u> | <u>30.2</u> | <u>31.2</u> | - | - |
| <u>19-Tongue</u> | <u>GenePrint</u> | <u>D21S11</u> | <u>30.2</u> | <u>31.2</u> | - | - |
| <u>20-Testis</u> | <u>GenePrint</u> | <u>D21S11</u> | <u>30.2</u> | <u>31.2</u> | - | - |
| <u>21-Thyroid</u> | <u>GenePrint</u> | <u>D21S11</u> | <u>30.2</u> | <u>31.2</u> | - | - |
| <u>22-Tibialis</u> | <u>GenePrint</u> | <u>D21S11</u> | <u>30.2</u> | <u>31.2</u> | - | - |
| <u>23-Visual Cortex</u> | <u>GenePrint</u> | <u>D21S11</u> | <u>30.2</u> | <u>31.2</u> | - | - |
| <u>01-Blood</u> | <u>GenePrint</u> | <u>D5S818</u> | <u>11</u> | <u>12</u> | - | - |
| <u>02-Brain Stem</u> | <u>GenePrint</u> | <u>D5S818</u> | <u>11</u> | <u>12</u> | - | - |
| <u>03-Cerebellum</u> | <u>GenePrint</u> | <u>D5S818</u> | <u>11</u> | <u>12</u> | - | - |
| <u>04-Colon</u> | <u>GenePrint</u> | <u>D5S818</u> | <u>11</u> | <u>12</u> | - | - |
| <u>05-CSF</u> | <u>GenePrint</u> | <u>D5S818</u> | <u>11</u> | <u>12</u> | - | - |
| <u>06-Gastrocnemius</u> | <u>GenePrint</u> | <u>D5S818</u> | <u>11</u> | <u>12</u> | - | - |
| <u>07-Heart</u> | <u>GenePrint</u> | <u>D5S818</u> | <u>11</u> | <u>12</u> | - | - |
| <u>08-Hippocampus</u> | <u>GenePrint</u> | <u>D5S818</u> | <u>11</u> | <u>12</u> | - | - |
| <u>09-Hypophysis</u> | <u>GenePrint</u> | <u>D5S818</u> | <u>11</u> | <u>12</u> | - | - |
| <u>10-Kidney</u> | <u>GenePrint</u> | <u>D5S818</u> | <u>11</u> | <u>12</u> | - | - |
| <u>11-Larynx</u> | <u>GenePrint</u> | <u>D5S818</u> | <u>11</u> | <u>12</u> | - | - |
| <u>12-Liver</u> | <u>GenePrint</u> | <u>D5S818</u> | <u>11</u> | <u>11</u> | - | - |
| <u>13-NMJ</u> | <u>GenePrint</u> | <u>D5S818</u> | <u>11</u> | <u>12</u> | - | - |
| <u>14-Pancreas</u> | <u>GenePrint</u> | <u>D5S818</u> | <u>11</u> | <u>12</u> | - | - |
| <u>15-Skin</u> | <u>GenePrint</u> | <u>D5S818</u> | <u>11</u> | <u>12</u> | - | - |
| <u>16-Spinal_Cord</u> | <u>GenePrint</u> | <u>D5S818</u> | <u>11</u> | <u>12</u> | - | - |
| <u>17-Stomach</u> | <u>GenePrint</u> | <u>D5S818</u> | <u>11</u> | <u>12</u> | - | - |
| <u>18-Striatum</u> | <u>GenePrint</u> | <u>D5S818</u> | <u>11</u> | <u>12</u> | - | - |
| <u>19-Tongue</u> | <u>GenePrint</u> | <u>D5S818</u> | <u>11</u> | <u>12</u> | - | - |
| <u>20-Testis</u> | <u>GenePrint</u> | <u>D5S818</u> | <u>11</u> | <u>12</u> | - | - |
| <u>21-Thyroid</u> | <u>GenePrint</u> | <u>D5S818</u> | <u>11</u> | <u>12</u> | - | - |
| <u>22-Tibialis</u> | <u>GenePrint</u> | <u>D5S818</u> | <u>11</u> | <u>12</u> | - | - |
| <u>23-Visual_Cortex</u> | <u>GenePrint</u> | <u>D5S818</u> | <u>11</u> | <u>12</u> | - | - |
| <u>01-Blood</u> | <u>GenePrint</u> | <u>D7S820</u> | <u>8</u> | <u>11</u> | - | - |
| <u>02-Brain Stem</u> | <u>GenePrint</u> | <u>D7S820</u> | <u>8</u> | <u>11</u> | - | - |
| <u>03-Cerebellum</u> | <u>GenePrint</u> | <u>D7S820</u> | <u>8</u> | <u>11</u> | - | - |
| <u>04-Colon</u> | <u>GenePrint</u> | <u>D7S820</u> | <u>8</u> | <u>11</u> | - | - |
| <u>05-CSF</u> | <u>GenePrint</u> | <u>D7S820</u> | <u>8</u> | <u>11</u> | - | - |
| <u>06-Gastrocnemius</u> | <u>GenePrint</u> | <u>D7S820</u> | <u>8</u> | <u>11</u> | - | - |
| <u>07-Heart</u> | <u>GenePrint</u> | <u>D7S820</u> | <u>8</u> | <u>11</u> | - | - |
| <u>08-Hippocampus</u> | <u>GenePrint</u> | <u>D7S820</u> | <u>8</u> | <u>11</u> | - | - |
| <u>09-Hypophysis</u> | <u>GenePrint</u> | <u>D7S820</u> | <u>8</u> | <u>11</u> | - | - |
| <u>10-Kidney</u> | <u>GenePrint</u> | <u>D7S820</u> | <u>8</u> | <u>11</u> | - | - |
| <u>11-Larynx</u> | <u>GenePrint</u> | <u>D7S820</u> | <u>8</u> | <u>11</u> | - | - |
| <u>12-Liver</u> | <u>GenePrint</u> | <u>D7S820</u> | <u>8</u> | <u>11</u> | <u>15</u> | - |
| <u>13-NMJ</u> | <u>GenePrint</u> | <u>D7S820</u> | <u>8</u> | <u>11</u> | - | - |
| <u>14-Pancreas</u> | <u>GenePrint</u> | <u>D7S820</u> | <u>8</u> | <u>11</u> | - | - |
| <u>15-Skin</u> | <u>GenePrint</u> | <u>D7S820</u> | <u>8</u> | <u>11</u> | - | - |
| <u>16-Spinal_Cord</u> | <u>GenePrint</u> | <u>D7S820</u> | <u>8</u> | <u>11</u> | - | - |
| <u>17-Stomach</u> | <u>GenePrint</u> | <u>D7S820</u> | <u>8</u> | <u>11</u> | - | - |
| <u>18-Striatum</u> | <u>GenePrint</u> | <u>D7S820</u> | <u>8</u> | <u>11</u> | - | - |
| <u>19-Tongue</u> | <u>GenePrint</u> | <u>D7S820</u> | <u>8</u> | <u>11</u> | - | - |
| <u>20-Testis</u> | <u>GenePrint</u> | <u>D7S820</u> | <u>8</u> | <u>11</u> | - | - |
| <u>21-Thyroid</u> | <u>GenePrint</u> | <u>D7S820</u> | <u>8</u> | <u>11</u> | - | - |
| <u>22-Tibialis</u> | <u>GenePrint</u> | <u>D7S820</u> | <u>8</u> | <u>11</u> | - | - |
| <u>23-Visual_Cortex</u> | <u>GenePrint</u> | <u>D7S820</u> | <u>8</u> | <u>11</u> | - | - |
| <u>01-Blood</u> | <u>GenePrint</u> | <u>TH01</u> | <u>6</u> | <u>9</u> | - | - |
| <u>02-Brain_Stem</u> | <u>GenePrint</u> | <u>TH01</u> | <u>6</u> | <u>6</u> | - | - |
| <u>03-Cerebellum</u> | <u>GenePrint</u> | <u>TH01</u> | <u>6</u> | <u>6</u> | - | - |
| <u>04-Colon</u> | <u>GenePrint</u> | <u>TH01</u> | <u>6</u> | <u>6</u> | - | - |
| <u>05-CSF</u> | <u>GenePrint</u> | <u>TH01</u> | <u>6</u> | <u>6</u> | - | - |
| <u>06-Gastrocnemius</u> | <u>GenePrint</u> | <u>TH01</u> | <u>6</u> | <u>6</u> | - | - |
| <u>07-Heart</u> | <u>GenePrint</u> | <u>TH01</u> | <u>6</u> | <u>6</u> | - | - |
| <u>08-Hippocampus</u> | <u>GenePrint</u> | <u>TH01</u> | <u>6</u> | <u>6</u> | - | - |
| <u>09-Hypophysis</u> | <u>GenePrint</u> | <u>TH01</u> | <u>6</u> | <u>6</u> | - | - |
| <u>10-Kidney</u> | <u>GenePrint</u> | <u>TH01</u> | <u>6</u> | <u>6</u> | - | - |
| <u>11-Larynx</u> | <u>GenePrint</u> | <u>TH01</u> | <u>6</u> | <u>6</u> | - | - |
| <u>12-Liver</u> | <u>GenePrint</u> | <u>TH01</u> | <u>6</u> | <u>9</u> | <u>10</u> | - |
| <u>13-NMJ</u> | <u>GenePrint</u> | <u>TH01</u> | <u>6</u> | <u>6</u> | - | - |
| <u>14-Pancreas</u> | <u>GenePrint</u> | <u>TH01</u> | <u>6</u> | <u>6</u> | - | - |
| <u>15-Skin</u> | <u>GenePrint</u> | <u>TH01</u> | <u>6</u> | <u>6</u> | - | - |
| <u>16-Spinal_Cord</u> | <u>GenePrint</u> | <u>TH01</u> | <u>6</u> | <u>6</u> | - | - |
| <u>17-Stomach</u> | <u>GenePrint</u> | <u>TH01</u> | <u>6</u> | <u>6</u> | - | - |
| <u>18-Striatum</u> | <u>GenePrint</u> | <u>TH01</u> | <u>6</u> | <u>6</u> | - | - |
| <u>19-Tongue</u> | <u>GenePrint</u> | <u>TH01</u> | <u>6</u> | <u>6</u> | - | - |
| <u>20-Testis</u> | <u>GenePrint</u> | <u>TH01</u> | <u>6</u> | <u>6</u> | - | - |
| <u>21-Thyroid</u> | <u>GenePrint</u> | <u>TH01</u> | <u>6</u> | <u>6</u> | - | - |
| <u>22-Tibialis</u> | <u>GenePrint</u> | <u>TH01</u> | <u>6</u> | <u>6</u> | - | - |
| <u>23-Visual_Cortex</u> | <u>GenePrint</u> | <u>TH01</u> | <u>6</u> | <u>6</u> | - | - |
| <u>01-Blood</u> | <u>GenePrint</u> | <u>TPOX</u> | <u>8</u> | <u>11</u> | - | - |
| <u>02-Brain_Stem</u> | <u>GenePrint</u> | <u>TPOX</u> | <u>8</u> | <u>11</u> | - | - |
| <u>03-Cerebellum</u> | <u>GenePrint</u> | <u>TPOX</u> | <u>8</u> | <u>11</u> | - | - |
| <u>04-Colon</u> | <u>GenePrint</u> | <u>TPOX</u> | <u>8</u> | <u>11</u> | - | - |
| <u>05-CSF</u> | <u>GenePrint</u> | <u>TPOX</u> | <u>8</u> | <u>11</u> | - | - |
| <u>06-Gastrocnemius</u> | <u>GenePrint</u> | <u>TPOX</u> | <u>8</u> | <u>11</u> | - | - |
| <u>07-Heart</u> | <u>GenePrint</u> | <u>TPOX</u> | <u>8</u> | <u>11</u> | - | - |
| <u>08-Hippocampus</u> | <u>GenePrint</u> | <u>TPOX</u> | <u>8</u> | <u>11</u> | - | - |
| <u>09-Hypophysis</u> | <u>GenePrint</u> | <u>TPOX</u> | <u>8</u> | <u>11</u> | - | - |
| <u>10-Kidney</u> | <u>GenePrint</u> | <u>TPOX</u> | <u>8</u> | <u>11</u> | - | - |
| <u>11-Larynx</u> | <u>GenePrint</u> | <u>TPOX</u> | <u>8</u> | <u>11</u> | - | - |
| <u>12-Liver</u> | <u>GenePrint</u> | <u>TPOX</u> | <u>8</u> | <u>11</u> | - | - |
| <u>13-NMJ</u> | <u>GenePrint</u> | <u>TPOX</u> | <u>8</u> | <u>11</u> | - | - |
| <u>14-Pancreas</u> | <u>GenePrint</u> | <u>TPOX</u> | <u>8</u> | <u>11</u> | - | - |
| <u>15-Skin</u> | <u>GenePrint</u> | <u>TPOX</u> | <u>8</u> | <u>11</u> | - | - |
| <u>16-Spinal_Cord</u> | <u>GenePrint</u> | <u>TPOX</u> | <u>8</u> | <u>11</u> | - | - |
| <u>17-Stomach</u> | <u>GenePrint</u> | <u>TPOX</u> | <u>8</u> | <u>11</u> | - | - |
| <u>18-Striatum</u> | <u>GenePrint</u> | <u>TPOX</u> | <u>8</u> | <u>11</u> | - | - |
| <u>19-Tongue</u> | <u>GenePrint</u> | <u>TPOX</u> | <u>8</u> | <u>11</u> | - | - |
| <u>20-Testis</u> | <u>GenePrint</u> | <u>TPOX</u> | <u>8</u> | <u>11</u> | - | - |
| <u>21-Thyroid</u> | <u>GenePrint</u> | <u>TPOX</u> | <u>8</u> | <u>11</u> | - | - |
| <u>22-Tibialis</u> | <u>GenePrint</u> | <u>TPOX</u> | <u>8</u> | <u>11</u> | - | - |
| <u>23-Visual_Cortex</u> | <u>GenePrint</u> | <u>TPOX</u> | <u>8</u> | <u>11</u> | - | - |
| <u>01-Blood</u> | <u>GenePrint</u> | <u>vWA</u> | <u>14</u> | <u>17</u> | - | - |
| <u>02-Brain_Stem</u> | <u>GenePrint</u> | <u>vWA</u> | <u>14</u> | <u>18</u> | - | - |
| <u>03-Cerebellum</u> | <u>GenePrint</u> | <u>vWA</u> | <u>14</u> | <u>18</u> | - | - |
| <u>04-Colon</u> | <u>GenePrint</u> | <u>vWA</u> | <u>14</u> | <u>18</u> | - | - |
| <u>05-CSF</u> | <u>GenePrint</u> | <u>vWA</u> | <u>14</u> | <u>18</u> | - | - |
| <u>06-Gastrocnemius</u> | <u>GenePrint</u> | <u>vWA</u> | <u>14</u> | <u>18</u> | - | - |
| <u>07-Heart</u> | <u>GenePrint</u> | <u>vWA</u> | <u>14</u> | <u>18</u> | - | - |
| <u>08-Hippocampus</u> | <u>GenePrint</u> | <u>vWA</u> | <u>14</u> | <u>18</u> | - | - |
| <u>09-Hypophysis</u> | <u>GenePrint</u> | <u>vWA</u> | <u>14</u> | <u>18</u> | - | - |
| <u>10-Kidney</u> | <u>GenePrint</u> | <u>vWA</u> | <u>14</u> | <u>18</u> | - | - |
| <u>11-Larynx</u> | <u>GenePrint</u> | <u>vWA</u> | <u>14</u> | <u>18</u> | - | - |
| <u>12-Liver</u> | <u>GenePrint</u> | <u>vWA</u> | <u>14</u> | <u>17</u> | <u>18</u> | - |
| <u>13-NMJ</u> | <u>GenePrint</u> | <u>vWA</u> | <u>14</u> | <u>18</u> | - | - |
| <u>14-Pancreas</u> | <u>GenePrint</u> | <u>vWA</u> | <u>14</u> | <u>18</u> | - | - |
| <u>15-Skin</u> | <u>GenePrint</u> | <u>vWA</u> | <u>14</u> | <u>18</u> | - | - |
| <u>16-Spinal_Cord</u> | <u>GenePrint</u> | <u>vWA</u> | <u>14</u> | <u>18</u> | - | - |
| <u>17-Stomach</u> | <u>GenePrint</u> | <u>vWA</u> | <u>14</u> | <u>18</u> | - | - |
| <u>18-Striatum</u> | <u>GenePrint</u> | <u>vWA</u> | <u>14</u> | <u>18</u> | - | - |
| <u>19-Tongue</u> | <u>GenePrint</u> | <u>vWA</u> | <u>14</u> | <u>18</u> | - | - |
| <u>20-Testis</u> | <u>GenePrint</u> | <u>vWA</u> | <u>14</u> | <u>18</u> | - | - |
| <u>21-Thyroid</u> | <u>GenePrint</u> | <u>vWA</u> | <u>14</u> | <u>18</u> | - | - |
| <u>22-Tibialis</u> | <u>GenePrint</u> | <u>vWA</u> | <u>14</u> | <u>18</u> | - | - |
| <u>23-Visual_Cortex</u> | <u>GenePrint</u> | <u>vWA</u> | <u>14</u> | <u>18</u> | - | - |

